# Asymmetric Signaling Across the Hierarchy of Cytoarchitecture within the Human Connectome

**DOI:** 10.1101/2022.05.13.491642

**Authors:** Linden Parkes, Jason Z Kim, Jennifer Stiso, Monica E Calkins, Matthew Cieslak, Raquel E Gur, Ruben C Gur, Tyler M Moore, Mathieu Ouellet, David R Roalf, Russell T Shinohara, Daniel H Wolf, Theodore D Satterthwaite, Dani S Bassett

**Affiliations:** Department of Bioengineering, School of Engineering & Applied Science, University of Pennsylvania, Philadelphia, PA, 19104 USA; Department of Psychiatry, Perelman School of Medicine, University of Pennsylvania, Philadelphia, PA 19104, USA; Lifespan Brain Institute, University of Pennsylvania & Children’s Hospital of Philadelphia, Philadelphia, USA; Center for Biomedical Image Computing and Analytics, Perelman School of Medicine, University of Pennsylvania, Philadelphia, PA 19104 USA; Department of Neurology, Perelman School of Medicine, Philadelphia, PA 19104 USA; Department of Radiology, Perelman School of Medicine, Philadelphia, PA 19104 USA; Department of Electrical & Systems Engineering, School of Engineering & Applied Science, University of Pennsylvania, Philadelphia, PA 19104 USA; Penn Statistics in Imaging and Visualization Center, Department of Biostatistics, Epidemiology, and Informatics, University of Pennsylvania, Perelman School of Medicine, Philadelphia, Pennsylvania; Department of Electrical & Systems Engineering, School of Engineering & Applied Science, University of Pennsylvania, Philadelphia, PA, 19104 USA; Department of Physics & Astronomy, College of Arts & Sciences, University of Pennsylvania, Philadelphia, PA, 19104 USA; Santa Fe Institute, Santa Fe, NM 87501 USA

## Abstract

Cortical variations in cytoarchitecture form a sensory-fugal axis that systematically shapes regional profiles of extrinsic connectivity. Additionally, this axis is thought to guide signal propagation and integration across the cortical hierarchy. While human neuroimaging work has shown that this axis constrains local properties of the human connectome, it remains unclear whether it also shapes the asymmetric signaling that arises from higher-order connectome topology. Here, we used network control theory to examine the amount of energy required to propagate dynamics across the sensory-fugal axis. Our results revealed an asymmetry in this energy indicating that bottom-up transitions were easier to complete compared to top-down transitions. Supporting analyses demonstrated that this asymmetry was underpinned by a connectome topology that is wired to support efficient bottom-up signaling. Finally, we found that this asymmetry correlated with changes in intrinsic neuronal timescales and lessened throughout youth. Our results show that cortical variation in cytoarchitecture may guide the formation of macroscopic connectome topology.

## INTRODUCTION

Multiple lines of evidence suggest that the brain’s extrinsic structural connectivity is predicted from its cytoarchitecture^1–4^. This *structural model* suggests that the degree to which two regions share similar cytoarchitectural features predicts the distribution of their laminar projections. Critically, inter-regional similarity in cytoarchitecture varies gradually across the cortex, creating a sensory-fugal (S-F) axis^5,6^ that predicts regions’ profiles of extrinsic connectivity to the rest of the brain. This gradient positions contiguous visual and sensorimotor cortex at one end and distributed heteromodal association and paralimbic cortices at the other, and is correlated with other macroscopic gradients of brain structure and function^7–15^. Together, these multi-modal gradients form a hierarchy of brain organization that is thought to govern extrinsic connectivity^3^ and support the efficient propagation and integration of signals across the cortex^16–18^. However, the extent to which cytoarchitecture’s governance over connectivity manifests in the topology of the macroscopic structural connectome remains a key open question. Here, we examine whether the S-F axis constrains signal propagation across macroscopic connectome topology.

Convergent evidence spanning the past three decades supports the premise that neuronal signaling is shaped and constrained by a globally ordered cortical hierarchy^16,17,19,20^. External stimuli arrive at functionally specialized sensory cortices before propagating up modality-specific hierarchies to then apex at association and paralimbic regions responsible for functional integration. This convergent bottom-up signal propagation is complemented by far-reaching modulatory top-down signals^21–24^ that operate on longer timescales^25^ and that bind incoming sensory signals together to update predictive inferences about our environment and to complete goal-directed action^26,27^. Critically, these cooperative patterns of bottom-up and top-down signaling, and the asymmetries between them^23^, may be underpinned by graded variations in cortical cytoarchitecture^2,3,28^. Specifically, regions’ cytoarchitecture robustly predicts their extrinsic connectivity profiles^1^, including the strength^29^, distance^29^, and layer origination and termination^29,30^ of feedforward and feedback projections^23^. Further, interregional similarity in cytoarchitecture follows a clear S-F axis^5,6^, suggesting that where a region is situated along the cortical hierarchy characterizes its bottom-up and top-down connectivity with the rest of the brain, and thus explains its capacity to support signal propagation across the hierarchy. Consistent with this notion, regional variation in the T1w/T2w ratio—a neuroimaging measurement that is thought to be a proxy of the S-F axis of cytoarchitecture^5,31^—correlates with regions’ intrinsic timescales of neuronal activity^10^, demonstrating that cytoarchitecture tracks the progressive lengthening of neuronal oscillations associated with hierarchical information integration^25^. Additionally, the S-F axis also correlates with regional weighted degree from diffusion-weighted structural networks^32^, demonstrating that cytoarchitecture tracks local properties of macroscopic connectome topology.

Regional variations to cytoarchitecture are, in part, rooted in neurodevelopment^1– 3,33–37^. Differences in the developmental timing of neurogenesis leads to highly eulaminate regions—such as the primary visual cortex—developing more slowly than agranular regions^38^, suggesting that prenatal development lays the foundation for the S-F axis. Once laid, the S-F axis scaffolds the formation of extrinsic feedforward and feedback connections that traverse up and down the hierarchy^3^. This connectivity formation also appears to track the S-F axis in a developmentally-staged manner, with synaptogenesis peaking earlier in lower-order primary visual areas than in higher-order frontal cortex^39– 41^. Furthermore, neuroimaging research shows that macroscopic proxies of the S-F axis (e.g., T1-weighted features), as well as structural connectivity, continue to change throughout postnatal development^15,42–48^, suggesting that the S-F axis continues to shape connectome topology.

As the above literature demonstrates, the processes that govern patterns of extrinsic connectivity across the cortex are encoded by regional variations in cytoarchitecture, and this regional variation provides a blueprint for the refinement of connectivity throughout development. However, the extent to which the topology of the structural connectome can be leveraged to model bottom-up and top-down signal propagation across the S-F axis remains unknown. The literature reviewed above leads us to four predictions. First, if differences in extrinsic projections encoded by cytoarchitecture are reflected in connectome topology^32^, then we should be able to model asymmetries between bottom-up and top-down signal propagation across the S-F axis in humans *in vivo*. Notably, recent work has shown that the topology of the undirected structural connectome generates spatially varied patterns of signal propagation^49^ and asymmetric signaling^50^, suggesting that such asymmetry may be assessable using non-invasive neuroimaging. Second, if asymmetric signal propagation is produced specifically by the cytoarchitectonic hierarchy, then asymmetries may not generalize to different views of the cortical hierarchy, such as those derived from patterns of functional connectivity^12^. Third, since signals propagating across the S-F axis will traverse through changing temporal receptive windows^10^, we expect asymmetries to correlate with differences in intrinsic neuronal timescales. Fourth, if signal propagation continues to be refined throughout development, then asymmetries should vary systemically as a function of age in youth.

To evaluate evidence for the above reasoning, we turned to the minimum control energy framework from Network Control Theory (NCT)^51,52^. Using a linear model of dynamics, NCT estimates the amount of input energy—delivered to a set of control nodes (brain regions)—that is required to drive the brain to transition between pairs of activity states. In this context, we consider binary states in which one set of regions are active while the rest of the brain is inactive. Here, we sought to estimate the transition energy associated with trans-hierarchical state transitions. We found that bottom-up state transitions were more efficient (required less energy) compared to top-down transitions. We also observed that the hierarchical distance separating brain states correlated with the size of these energy asymmetries, suggesting that states with different underlying cytoarchitecture display the most pronounced asymmetries. In addition to these primary findings, we examined (i) whether our findings generalized to the principal gradient of functional connectivity^12^; (ii) whether our transition energies correlated with between-state differences in intrinsic timescales; (iii) whether brain regions’ position along the S-F axis explained their role in facilitating state transitions; and (iv) whether energy asymmetries correlated with age in a developing sample of youths. Our work extends the field’s understanding of connectome topology by showing that the neuroanatomical processes that give rise to extrinsic connectivity constrain the directional flow of macroscopic dynamics over the cortex.

## RESULTS

### Mapping trans-hierarchical state transitions

We characterized the energy required to complete trans-hierarchical state transitions. Here, we set our brain states to actuate patches of cortex with relatively homogenous profiles of cytoarchitecture. Briefly, we defined brain states by splitting the S-F axis of cytoarchitectonic similarity^5,6^ into *k* equally sized non-overlapping groups of regions that spanned the gradient (**Fig. 1A**; see Methods). Then, using a group-averaged structural connectome taken from the Philadelphia Neurodevelopmental Cohort^53^ (see Methods), we modeled the transition energy between all *k* pairs of brain states, generating a *k* × *k* matrix of energy values, *T*_*E*_ (**Fig. 1B, C**). Critically, the hierarchically ordered nature of our brain states meant that bottom-up transition energies were naturally stored in the upper triangle of *T*_*E*_ while top-down transition energies were stored in the lower triangle. We computed energy asymmetries by subtracting top-down energy from bottom-up energy (**Fig. 1D**; 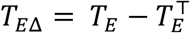). In the upper triangle of *T*_*E*Δ_, positive values indicate bottom-up energy being greater than top-down energy whereas negative values indicate bottom-up energy being lower than top-down energy.

**Figure 1.**
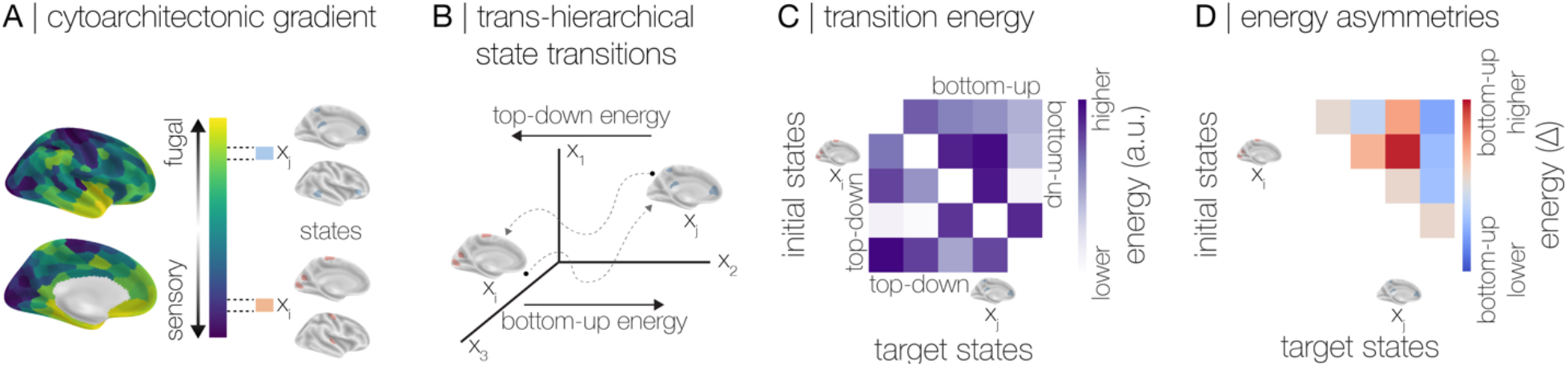
Estimating trans-hierarchical signal propagation. Using the Schaefer atlas, we sampled 20 non-overlapping groups of regions (*n*=10 per state) traversing up the S-F gradient of cytoarchitectonic similarity^6^. These groups formed brain states spanning the cortical hierarchy. By definition, regions within each state had similar profiles of cyoarchitecture. Accordingly, pairs of states separated by long hierarchical distances have different underlying cytoarchitecture. **A**, An example pair of brain states (*x*_*i*_, *x*_*j*_) at different locations along the cytoarchitectonic hierarchy. **B**, For a given pair of states (*x*_*i*_, *x*_*j*_), we calculated the minimum control energy (*E*) required to complete the transition from *x*_*i*_ to *x*_*j*_ and from *x*_*j*_ to *x*_*i*_. **C**, Minimum control energy between all pairs of states was assembled into a transition energy matrix, *T*_*E*_. Owing to the ordered nature of our brain states, transition energies were trivially grouped into bottom-up (transitions moving up the hierarchy; *T*_*E*_, upper triangle) and top-down (transitions moving down the hierarchy; *T*_*E*_, lower triangle). **D**, Given this grouping, we subtracted top-down energy from bottom-up energy to create an energy asymmetry matrix (*T*_*E*Δ_). In the upper triangle of this asymmetry matrix, positive values represented state transitions where bottom-up energy was higher than top-down energy whereas negative values represented the opposite. Note that, apart from the sign of the Δ value, *T*_*E*Δ_ is symmetric; hence, all analyses of asymmetries focused on the upper triangle of this matrix.

We found that bottom-up energy was significantly lower than top-down energy (**Fig. 2A**; *t*=5.94, *p*=1×10^−8^), demonstrating that state transitions moving up the cytoarchitectonic S-F axis required less energy (i.e., were easier to complete) compared to those moving down the same axis. Furthermore, in support of our hypothesis, we found that the hierarchical distance between brain states was negatively correlated with *T*_*E*Δ_ (**Fig. 2B**, left). That is, as states’ cytoarchitecture became more dissimilar from one another (greater distance), energy asymmetries became more negative. Thus, asymmetries between bottom-up and top-down transition energies were largest when brain states had differing cytoarchitecture, with bottom-up transitions becoming progressively easier to complete than top-down. We found convergent results when we re-ran analyses on a single hemisphere, thereby excluding inter-hemispheric connections (bottom-up energy *versus* top-down, *t*=3.31, *p*=2×10^−3^; correlation with hierarchical distance, *ρ*=-0.30, *p*_*parametric*_=5×10^−2^).

**Figure 2.**
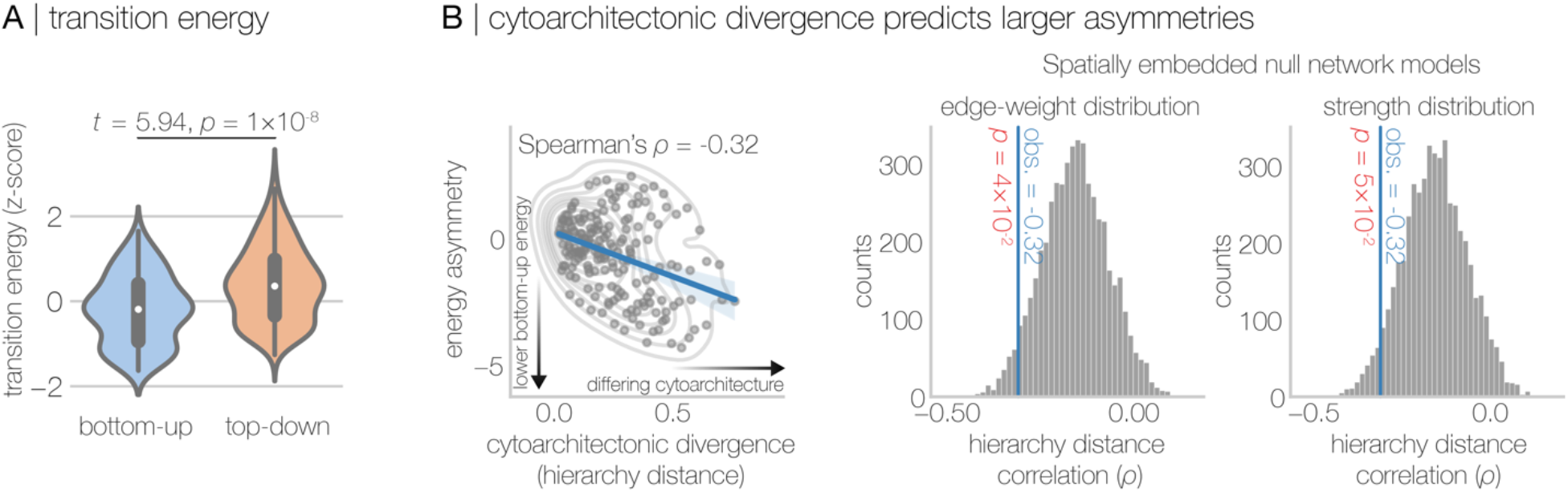
The topology of the structural connectome is sensitive to asymmetries between top-down and bottom-up signal propagation across the sensory-fugal axis of cytoarchitecture. **A**, Bottom-up energy was significantly lower than top-down energy, demonstrating that bottom-up state transitions were easier for our network control model to complete. **B**, The distance along the cytoarchitectonic gradient separating initial and target states was negatively correlated with energy asymmetry, demonstrating that high cytoarchitectonic dissimilarity between states was linked to greater negative energy asymmetries (left). This finding shows that when cytoarchitecture differs between brain states, bottom-up transitions required lower energy to complete compared to their top-down counterparts. This correlation with hierarchy distance was larger than expected under a pair of null network models (right), including one that preserved the spatial embedding and the edge weight distribution of the network and another that preserved the spatial embedding and the strength distribution. This observation suggests that this hierarchy distance effect may be supported by higher-order topology of the structural connectome.

Next, to examine this distance effect’s dependence on topology, we recomputed *T*_*E*Δ_ under two null network models. Specifically, we randomly rewired the underlying group-averaged structural connectome 5,000 times using a spatially embedded permutation model that preserved either the edge distribution or the strength distribution of the network^54^ (see Methods). Then, for every rewired connectome, we re-estimated *T*_*E*Δ_ as well as the correlation with hierarchical distance. We found that the observed correlation was stronger than expected under both null distributions (**Fig. 2B**, right). This result demonstrates that the negative correlation between hierarchical distance and *T*_*E*Δ_ was not simply explained by a combination of the network’s spatial embedding and edge (or strength) distribution, suggesting instead that this effect may be specifically explained by variations in cytoarchitectonic profiles.

To test whether our results were specific to the S-F axis of cytoarchitecture, we repeated all of the above analyses using the principal gradient of functional connectivity^12^ to define brain states (**Fig. S1**). This is a relatively strong test of specificity as the gradient of cytoarchitecture and the gradient of functional connectivity were correlated (*r*=0.594). Using the functional connectivity gradient, we found that bottom-up and top-down transition energies did not differ significantly (**Fig. S1A**; *t*=1.147, *p*=0.142). Additionally, we observed a relatively weak positive correlation between hierarchical distance and *T*_*E*Δ_ that was not larger than expected under our null network models (**Fig. S1B**; *ρ*=0.11, *p*_edge_=0.902, *p*_strength_=0.843). This result demonstrates that asymmetries between bottom-up and top-down transition energies were specific to the S-F axis of cytoarchitecture. This lack of energy asymmetry for the functional gradient may be explained by the fact that the two axes diverge at their apex^5,55^; the top of the S-F axis comprises paralimbic regions while the top of the functional connectivity axis comprises transmodal cortex. Previous work has suggested that this (relative) untethering of functional connectivity from cytoarchitectonic constraints may support the functional diversity of the transmodal cortex^5^. This untethering is also consistent with evidence that macroscopic structural and functional connectivity are relatively uncoupled in transmodal cortex compared to unimodal cortex^56,57^. Thus, together with past literature, our findings converge on the idea that while cytoarchitecture and structural connectivity are tightly intertwined, functional connectivity departs from both in a spatially patterned way.

### The gradient of cytoarchitecture constrains the flow of activity over the cortex

The above results demonstrate that the energy asymmetries associated with trans-hierarchical state transitions may be a consequence of more than brain regions’ spatial embedding and the distribution (or strength) of their direct links. Specifically, this evidence suggests that the entire pattern of connectome topology, not just direct connections, may be optimized to propagate activity up the cytoarchitectonic gradient more efficiently compared to down, in turn enabling more efficient completion of bottom-up state transitions. To probe this possibility further, we examined whether the flow of uncontrolled activity followed the S-F axis as it spread throughout the cortex over time (**Fig. 3A**; see Methods). Briefly, seeding from each brain state, we examined the spread of natural dynamics across the whole brain as they unfolded over time (i.e., over a series of time steps; see Methods). Intuitively, this amounted to re-simulating our dynamical model for each initial state in the absence of both a target state and a control set. In a pair of analyses described below, we used this approach to show that the topology of the connectome may be optimized to propagate activity up the hierarchy.

**Figure 3.**
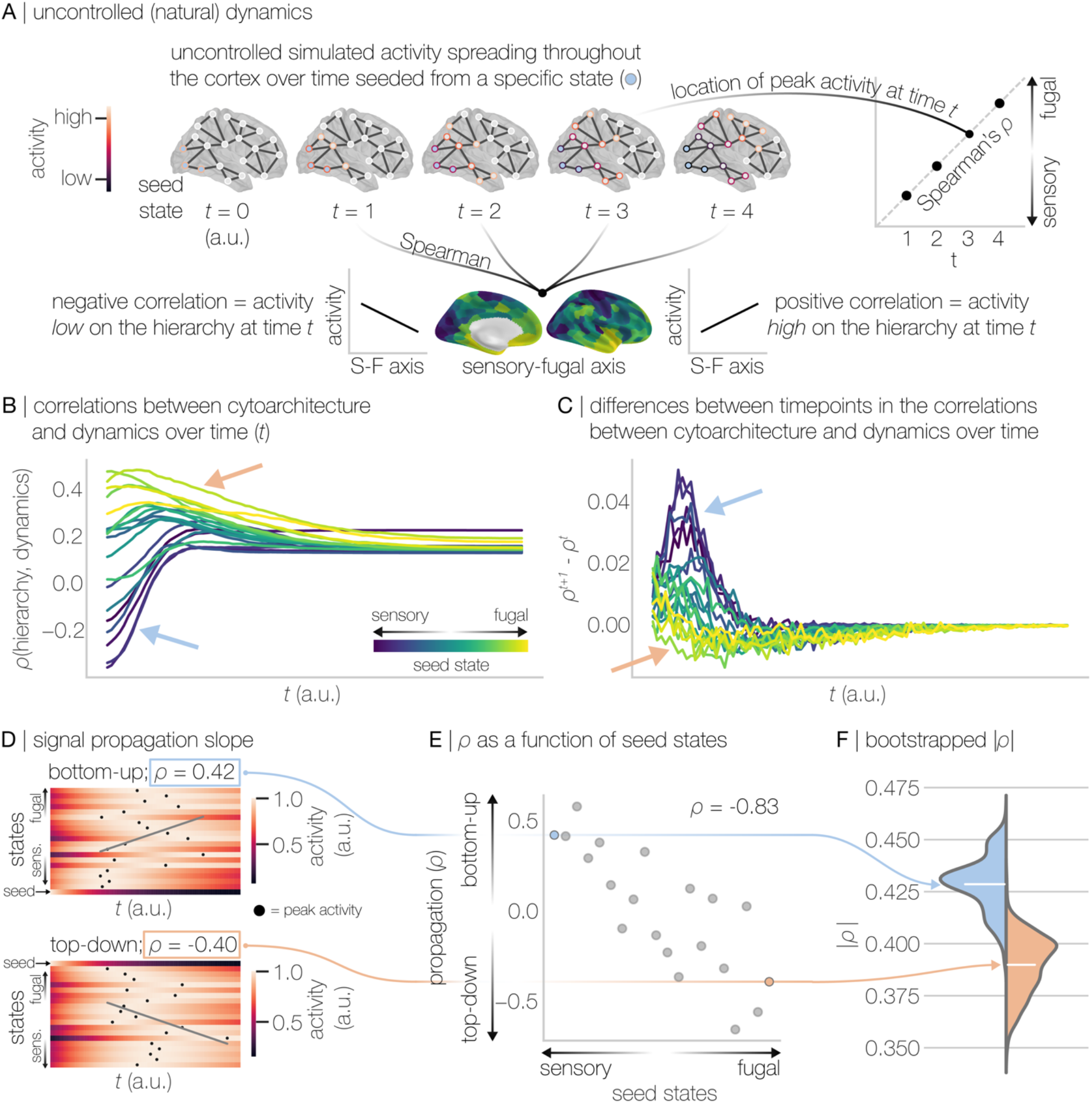
Uncontrolled dynamics preferentially flow up the cortical gradient of cytoarchitecture. **A**, We simulated the spread of uncontrolled dynamics seeded from each of our cytoarchitectonic brain states and tracked the activity as it unfolded over time and spread throughout the cortex. For a given seed state, we performed two analyses. First, we quantified the Spearman rank correlation between the sensory-fugal axis of cytoarchitecture and the pattern of simulated activity at time *t* (results in panel **B**) as well as the difference in correlations between adjacent timepoints (results in panel **C**). Second, we examined when activity peaked within each of the other cytoarchitectonic brain states (results shown in panels **D, E**, and **F**). We quantified this peak effect by calculating the Spearman rank correlation between *t* and the location (i.e., state) of peak activity. **B**, Correlations between cytoarchitecture and simulated activity seeded from each brain state as a function of time. **C**, Differences in correlations between neighboring timepoints as a function of time. **D**, Correlation between the location of peak activity and time. Activity seeded from the bottom of the S-F axis traversed up the gradient (top) whereas activity seeded from the top of the S-F axis traversed down the gradient (bottom). Black markers denote the timepoints when activity peaked within each brain state as it traversed the S-F axis. **E**, These propagating patterns of simulated activity were summarized using Spearman rank correlations separately for each seeded brain state. **F**, Finally, under 1,000 bootstrapped samples of the group-averaged connectome, we found that the magnitude of the bottom-up Spearman correlation (seeded from the bottom of the S-F axis) was significantly larger than the magnitude of the top-down Spearman correlation (seeded from the top of the S-F axis). Collectively, these results suggest that uncontrolled dynamics spread more readily across the S-F axis in the bottom-up direction than top-down.

First, we correlated the pattern of activity observed at each time step (*t*, arbitrary units) with the S-F axis (**Fig. 3B**). Here, for a given time step, negative correlations indicated that brain activity was higher at the bottom of the hierarchy than at the top, while positive correlations indicated the opposite. **Fig. 3B** shows that states lower on the hierarchy tend to show negative correlations between the S-F axis and early activity propagation (**Fig. 3B**, blue arrow), while states higher on the hierarchy tend to show positive correlations (**Fig. 3B**, peach arrow). Expectedly, this pattern demonstrates that early signal propagation tends to activate regions near to a given state’s location on the hierarchy. That is, activity propagating from low positions on the hierarchy reaches other low-hierarchy regions first, driving a negative correlation, while activity propagating from high on the hierarchy reaches other high-hierarchy regions first, driving a positive correlation. Critically, **Fig. 3B** also shows that the negative correlations low on the hierarchy diminish (i.e., become less negative) more quickly compared to the positive correlations for the high-hierarchy states. This effect is quantified and recapitulated in **Fig. 3C**, which shows the differences in correlations between neighboring time points (ρ ^*t*−1^ − ρ ^*t*^). Specifically, we found that differences in correlations between timepoints were greater when activity was seeded from the bottom of the hierarchy (**Fig. 3C**, blue arrow) compared to the top (**Fig. 3C**, peach arrow). Together, these results suggest that activity propagates more readily in the bottom-up direction than in the top-down direction.

Second, we sought to stringently assess this apparent difference between bottom-up and top-down propagation efficiency. For each seeded brain state, we identified the point in time when simulated activity peaked within each of the other brain states (see Methods). Then, to quantify the slope of this activity propagation, we calculated the Spearman correlation between these peak time points and states’ position on the hierarchy (**Fig. 3D, E**). Thus, this analysis quantified the extent to which activity spreading from each brain state peaked within the remaining brain states in an ordered fashion. Lastly, we regenerated the group averaged connectome 500 times using bootstrapping, which allowed us to test for differences in these slopes using confidence intervals (see Methods). In doing so, we found that the correlation quantifying bottom-up propagation from the lowest position on the S-F axis (mean |*ρ*|=0.4286; 95% CI=[0.4278, 0.4294]) was significantly larger than the correlation quantifying top-down propagation from the topmost position (mean |*ρ*|=0.3899; 95% CI=[0.3890, 0.3907]) (**Fig. 3F**). This result provides evidence that waves of natural dynamics flowing up the S-F axis tend to traverse the hierarchy more readily than their top-down counterparts. Collectively, the results presented in **Fig. 3** suggest that cytoarchitecture may constrain the topology of the network to enable more efficient bottom-up flow of information. Furthermore, these results are consistent with our observation of lower bottom-up energy compared to top-down (see **Fig. 2**); a topology that is organized to facilitate bottom-up activity flow will require less energy to complete controlled bottom-up state transitions compared to top-down.

### Energy asymmetries in trans-hierarchical state transitions are correlated with differences in intrinsic timescales and asymmetries in effective connectivity

Our observations thus far are consistent with the notion that regional cytoarchitectonic similarity influences the difference between bottom-up and top-down signal propagation across the cortical hierarchy. Specifically, our results suggest that how patterns of brain activity spread across the hierarchy varies as a function of the direction of flow. However, the results presented thus far were only derived from linear dynamics simulated upon the structural connectome. We reasoned that if our results for simulated dynamics were neurobiologically meaningful, then we would observe two findings.

First, we expected that energy asymmetries would correlate with changes in the intrinsic neuronal timescales of our brain states. Specifically, we predicted that transitions where bottom-up energy was lower than top-down would correspond to a lengthening of neuronal timescales between the initial and target states. In turn, this finding would suggest that the topology of the structural connectome is wired to support the integration of information that is thought be occurring as activity traverses up the hierarchy. Second, we expected that energy asymmetries would be consistent with asymmetries derived from dynamical models trained on functional neuroimaging data. To test the former prediction, we used open-access human electrocorticography (ECoG) data^58,59^ to index regions’ intrinsic timescales. Specifically, following Gao *et al*.^10^, we quantified timescales using the time constant (τ) of an exponential decay function fitted to the autocorrelation function of the ECoG timeseries (**Fig. 4A**; see Methods). Larger τ values correspond to longer (slower) fluctuations in a region’s intrinsic timescales. Subsequently, we averaged τ within each of our brain states and then subtracted mean τ between pairs of brain states (τ_Δ_). Thus, positive τ_Δ_ represented larger τ in state *j* compared to state *i*. Finally, we correlated *T*_*E*Δ_ with τ_Δ_ and found that they were negatively correlated (**Fig. 4B**; *r*=-0.34, *p*=1×10^−6^). This result indicates that state transitions where bottom-up energy is lower than top-down (i.e., negative *T*_*E*Δ_) are also characterized by an increase in τ (i.e., positive τ_Δ_) going from state *i* to state *j* and *vice versa*. Thus, state transitions that are (relatively) easy to complete are coincident with a lengthening of the timescales of resting-state electrophysiological fluctuations.

**Figure 4.**
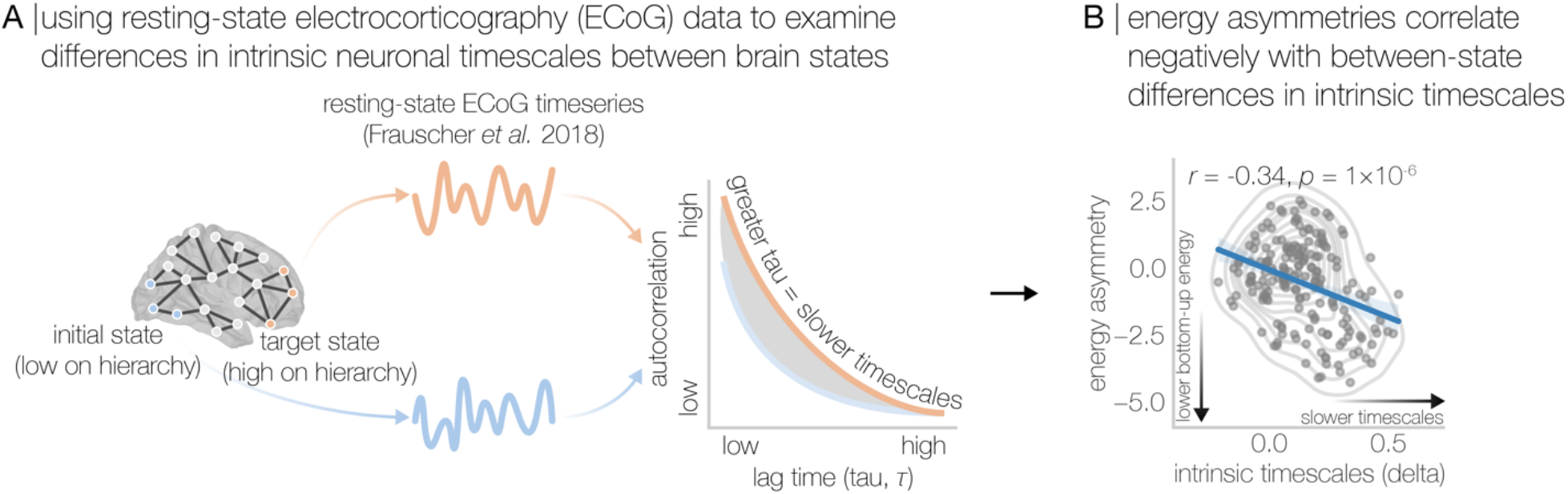
Energy asymmetries correlate with differences between brain states’ intrinsic neuronal timescales. **A**, We used resting-state electrocorticography data to examine differences between brain states’ intrinsic neuronal timescales (as per methods described in Gao *et al*.^10^) between our cytoarchitectonic brain states. **B**, Energy asymmetries between brain states were negatively correlated with differences between brain states’ intrinsic timescales. This result shows that state transitions where bottom-up energy is lower than top-down (negative energy asymmetry) are also characterized by a slowing of intrinsic timescales going from state *i* to state *j* and *vice versa*.

Next, we turned from an evaluation of differences between brain states’ intrinsic neuronal timescales to an evaluation of asymmetries derived from effective connectivity. Specifically, we computed the effective connectivity (EC) between brain states using a spectral version of dynamic causal modeling^60,61^ applied to participants’ resting-state functional magnetic resonance imaging (rs-fMRI) data (see Methods). We subsequently computed EC asymmetries by subtracting top-down EC from bottom-up EC (*EC*_Δ_ = |*EC*| − |*EC*|^⊤^). We found that *T*_*E*Δ_ was positively correlated with *EC*_Δ_ (**Fig. S2**; *r*=0.24, *p*=1×10^−3^), indicating that for state transitions where bottom-up energy was lower than top-down the same was true for EC and *vice versa*. This result extends prior work^50^ by demonstrating that the topology of the undirected structural connectome supports directed signal propagation along the cortical gradient of cytoarchitectonic similarity.

### Optimized control weights increase energy asymmetries and track the sensory-fugal axis of cytoarchitecture

The preceding sections demonstrated that brain network topology may be optimized to facilitate more efficient bottom-up trans-hierarchical state transitions compared to top-down, and that this effect (i) is not better explained by spatial embedding or lower-order topology, (ii) is specific to cytoarchitecture, and (iii) is consistent with asymmetries in intrinsic timescales and effective connectivity. Next, we sought to understand whether regions’ position along the hierarchy informed their capacity to facilitate trans-hierarchical state transitions. To achieve this goal, we optimized state transitions by introducing a variable set of control weights that minimized transition energy. These weights change the relative influence of control assigned to different brain regions, rather than assuming that all regions have equal influence. We predicted that optimized weights would maximize energy asymmetries and track the S-F axis. We tested these predictions in sequence, starting with the prediction that optimized weights would maximize energy asymmetries. For each state transition, we systematically perturbed the system to generate a set of control weights that minimized transition energy (**Fig. 5A**; see Methods). Then, upon these minimized transition energies, we recomputed *T*_*E*Δ_. Similar to our primary analysis using uniform control weights (see **Fig. 2A**), we found that bottom-up energy was significantly lower than top-down when using optimized control weights (**Fig. 5B**; *t*=6.46, *p*=9×10^−10^). However, this asymmetry in energies for optimized weights was larger compared to uniform control weights (*t*=6.46 in **Fig. 5B** versus *t*=5.94 in **Fig. 2A**). This finding is consistent with our first prediction.

**Figure 5.**
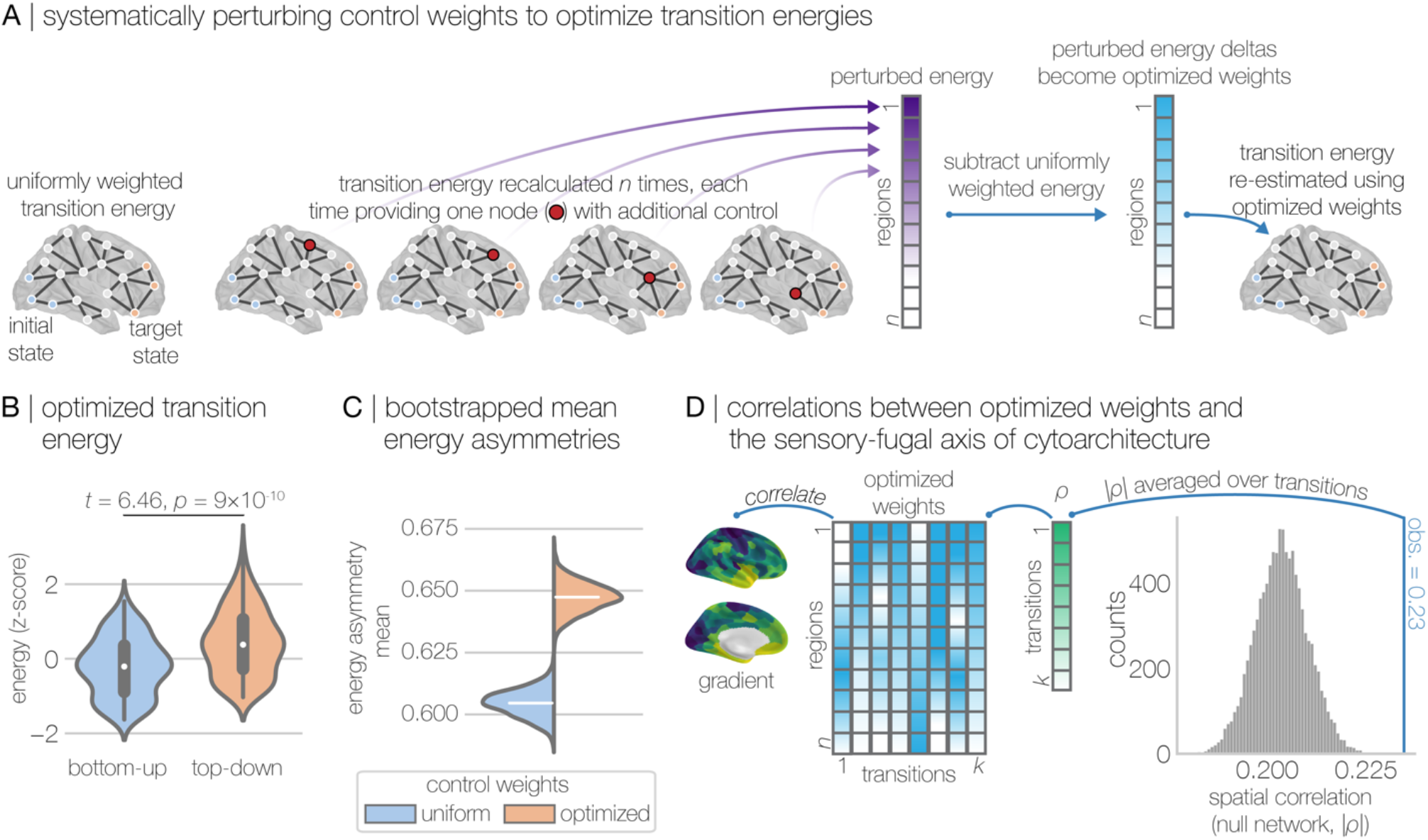
Optimized control weights maximize energy asymmetries and track the cortical gradient of cytoarchitecture. **A**, For each trans-hierarchical state transition, we adopted the following procedure to generate optimized control weights that minimized transition energy. First, for a given state transition, we calculated uniformly weighted transition energy; nodes of the system were provided the same degree of control over system dynamics. Note, results for uniformly weighted transition energy have been reported in all figures prior to this one. Second, we re-estimated the transition energy *n* times, each time providing one node with additional control over the system. This approach generated a vector of perturbed transition energies (purple vector). Third, we subtracted the uniformly weighted energy from each of the perturbed energies to generate a vector of perturbed energy deltas (blue vector), the magnitude of which encoded regions’ importance to the state transition. Fourth, we re-estimated transition energy one more time using the perturbed energy deltas as optimized control weights. **B**, Transition energies estimated using optimized control weights. Consistent with results observed for uniform control weights (see **Fig. 2**), here using optimized control weights we found that bottom-up energy was significantly lower than top-down energy. **C**, Mean energy asymmetries (|*T*_*E*Δ_|) for uniform (blue) compared to optimized (peach) control weights under 500 bootstraps (see Methods). Mean |*T*_*E*Δ_| was unambiguously larger for optimized control weights (mean |*T*_*E*Δ_|=0.6474; 95% CI=[0.6470, 0.6478]) compared to uniform control weights (mean |*T*_*E*Δ_|=0.6046; 95% CI=[0.6043, 0.6049]). **D**, Spatial correlations between optimized control weights and the S-F axis of cytoarchitecture averaged over state transitions. This average spatial correlation was larger than expected under a null network model that preserved spatial embedding and the strength distribution of the nodes.

To assess whether this difference in asymmetry sizes was robust, we computed mean *T*_*E*Δ_ for both uniform and optimized control weights under 500 bootstrapped samples (see Methods). **Fig. 5C** shows that mean *T*_*E*Δ_ was unambiguously larger for optimized control weights (mean |*T*_*E*Δ_|=0.6474; 95% CI=[0.6470, 0.6478]) compared to uniform control weights (mean |*T*_*E*Δ_|=0.6046; 95% CI=[0.6043, 0.6049]), demonstrating that the former yielded significantly larger energy asymmetries. Note that optimized weights were only designed to minimize transition energy, including both bottom-up and top-down energies. Thus, this observed increase in mean |*T*_*E*Δ_| suggests that our optimized control weights minimized bottom-up energy to a greater extent than top-down.

Next, we turned to our second prediction, and examined the correlation between optimized control weights and the S-F axis (see Methods). Here, any observed correlation implies that optimizing control weights uncovers a spatial mode of control variation that tracks the gradient of cytoarchitecture. When averaged across state transitions, we found that the correlations between optimized control weights and cytoarchitecture were |ρ|=0.23±0.14 (this was true at the level of specific state transitions as well; see **Fig. S3**). Note that we examine and report absolute correlations here as we are interested in whether optimized weights couple to the gradient generally (signed correlations are reported in **Fig. S3**). Further, this correlation was significantly larger than expected under our spatially informed strength preserving null network model (**Fig. 5D**; and see **Fig. S3**). Together, these results illustrate that a region’s position along the S-F axis explains its role in facilitating trans-hierarchical state transitions, and that imbuing our model with knowledge of these roles optimizes the efficiency of bottom-up signal propagation across the hierarchy.

### Asymmetries in trans-hierarchical state transitions are refined throughout development

Having illustrated that a region’s position along the S-F axis explains its role in facilitating state transitions, we next sought to characterize the developmental trajectories of transition energies. Based on previous literature, we expected that ongoing developmental refinement of structural connectivity would result in age-related changes to bottom-up and top-down energy. To test this expectation, we estimated the correlation between participant-specific transition energies and age, while controlling for sex, total brain volume, and in-scanner motion (**Fig. 6A**). Here, energy was estimated using participant-specific optimized weights (see Methods and previous section). On average, we found that age correlated negatively with both bottom-up (**Fig. 6B**) and top-down energy (**Fig. 6C**), with the latter effect being stronger. These results illustrate that the energy asymmetry we observe in our data (see **Fig. 2A**) may weaken as a function of age. In turn, neurodevelopmental refinement of the connectome may involve converging toward a balance between bottom-up and top-down signal propagation. We also found that the transition-level age effects were negatively correlated with *T*_*E*Δ_ taken from the group-average connectome (**Fig. 6D**). This result demonstrates that state transitions with stronger energy asymmetries also showed the strongest age effects. Lastly, using a cross-validated penalized regression model (**Fig. 6E**; see Methods), we found that energy asymmetries were able to robustly predict participants’ age in out-of-sample testing (**Fig. 6F**; see also **Fig. S4** which shows that optimized energies better predicted participants’ age compared to non-optimized energies derived from uniform control weights). Consistent with our expectations, these results show that development plays a critical role in refining trans-hierarchical transition energies, and that this refinement is concentrated in state transitions with divergent cytoarchitecture.

**Figure 6.**
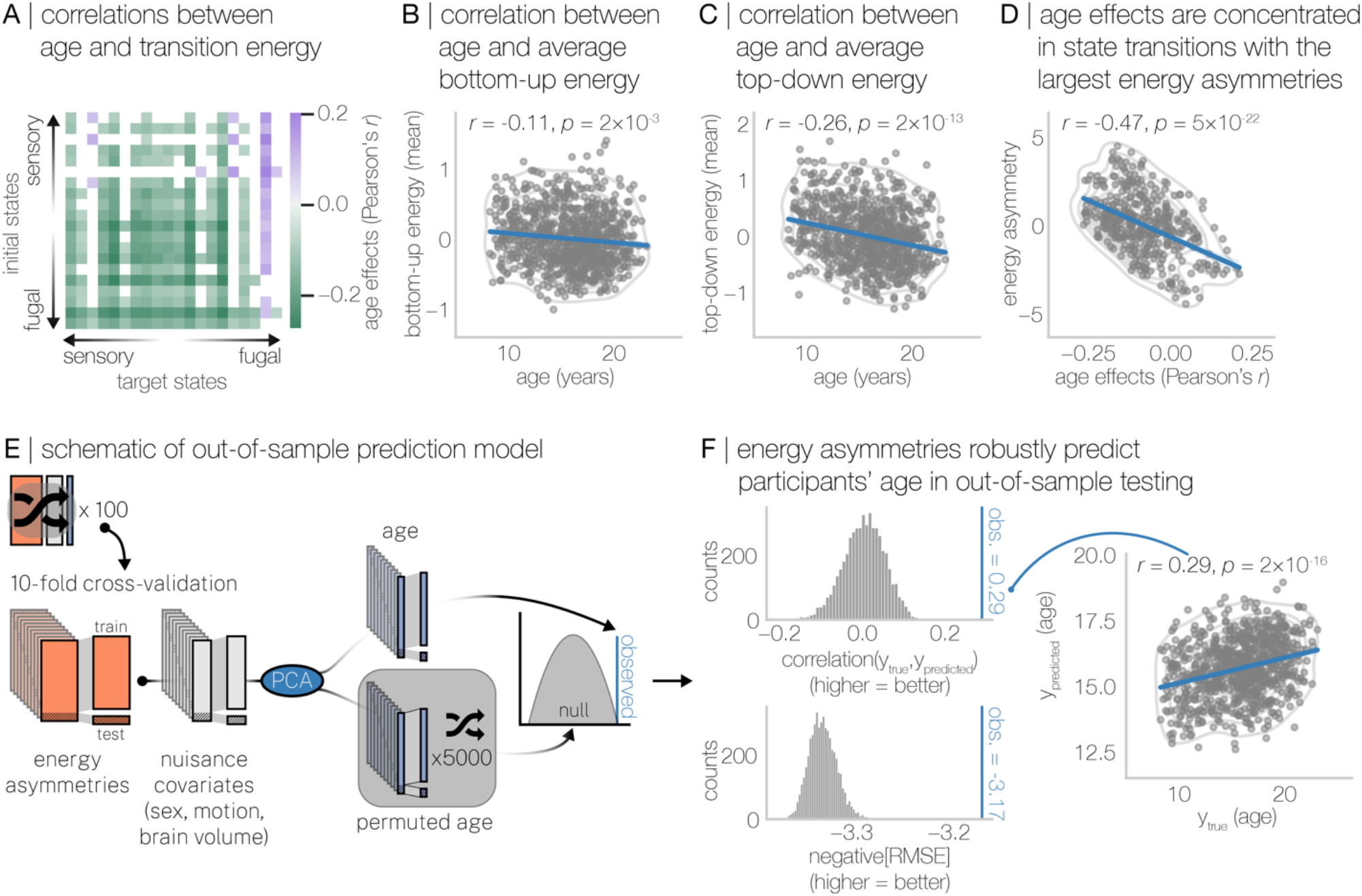
Energy asymmetries in trans-hierarchical state transitions vary systematically over development. We estimated correlations between age and trans-hierarchical transition energy in 793 individuals, while controlling for sex, total brain volume, and in-scanner motion (see Methods). **A**, Correlations between age and transition energy for all state transitions. We observed widespread negative correlations between age and transition energy, suggesting that state transitions became easier to complete as individuals got older. **B**, Correlation between age and average bottom-up energy. **C**, Correlation between age and average top-down energy. We found that while both bottom-up and top-down energy reduced as a function of age, the age effect for top-down energy was larger than that observed for bottom-up. This result suggests that the energy asymmetry between bottom-up and top-down closed throughout youth. **D**, Correlation between state level age effects (from **A**) and the *T*_*E*Δ_ derived from the group-averaged structural connectome (see **Fig. 2**). We found that the size of the age effects (Pearson’s *r*) was negatively correlated with *T*_*E*Δ_, demonstrating that the strongest age effects were concentrated in the state transitions with the largest energy asymmetries. **E**, Schematic illustration of a cross-validated regression model that was used to assess out-of-sample prediction of participants’ age. **F**, Results from out-of-sample prediction of participants’ age. Energy asymmetries robustly predicted participants’ age in out-of-sample testing when scored using both the correlation between true and predicted *y* (top, left) and negative root mean square error (bottom, left). Note, these prediction effects were replicated when using both a higher resolution version of our parcellation that included 400 parcels (Schaefer 400; correlation(y_true_, y_predicted_)=0.32; negative[RMSE]=-3.14) as well as a 360-parcel multi-modal parcellation developed in the Human Connectome Project (correlation(y_true_, y_predicted_)=0.31; negative[RMSE]=-3.14). Taken together, these results show clearly that asymmetries in trans-hierarchical signal propagation and neurodevelopment are intimately intertwined.

## DISCUSSION

Here, we investigated how the relationship between cytoarchitecture and connectivity constrains the dynamics engendered by the structural connectome. Using NCT^51,52,62,63^, we modeled the amount of control energy that was required to propagate linear dynamics up and down the S-F axis of cytoarchitecture. We reported several key findings. First, we found that the energy required to complete bottom-up state transitions was lower compared to their top-down counterparts, indicating that bottom-up transitions were easier for our model to complete. Additionally, through a combination of null network models and analyses of uncontrolled dynamics, we found that this energy asymmetry was underpinned by a network topology that is wired to enable efficient bottom-up signaling across the cortical hierarchy. Second, we found that energy asymmetries correlated with differences in intrinsic neuronal timescales estimated from ECoG as well as asymmetries in effective connectivity estimated from resting-state fMRI. The former finding demonstrates that efficient bottom-up signaling across the structural connectome is coincident with a lengthening of regions’ temporal receptive windows, while the latter shows that our model of dynamics is consistent with those drawn from functional data. Third, we found that regions’ position along the S-F axis was correlated with their importance in facilitating state transitions, demonstrating that the spatial modes of control embedded in our model were coupled to the cortical hierarchy. Finally, we found that bottom-up and top-down energy decreased as a function of age in a sample of developing youths. Notably, while age correlated negatively with both bottom-up and top-down energy, effects were more pronounced for top-down—indicating that energy asymmetries tended to lessen as a function of development in our sample—and reductions were strongest for the state transitions with the largest asymmetries. Overall, our results demonstrate that the higher-order topology of the human connectome may be wired to support asymmetric signaling across the cortical hierarchy, and that this signaling is rooted in the spatial patterning of cytoarchitecture that is itself guiding the ongoing refinement of connectivity throughout youth.

### Cytoarchitecture shapes the connectome

Understanding how cytoarchitecture shapes connectivity is a central goal of neuroscience^64^. In humans, recent research has shown a clear link between cortical cytoarchitecture and local properties of structural connectivity^32,65,66^. Using graph theory, Wei *et al*.^32^ found that several indices of regions’ local network importance correlated moderately with regions’ cytoarchitectonic similarity to the rest of the brain, demonstrating that regions with similar cytoarchitecture were more strongly, and more globally, connected to the rest of the network. Paquola *et al*.^66^ defined a regional embedding space that fused together edge-level structural connectivity, geodesic distance, and cytoarchitectonic similarity^5^. Paquola *et al*.^66^ found that their wiring diagram explained variance in regional externopyramidization—which tracks the laminar origin of neuronal projections^64^—supporting the notion that variance in the laminar origin of feedback and feedforward connections is intertwined with variance in cytoarchitecture and macroscopic connectivity. Our findings extend these prior studies by showing that cytoarchitecture shapes not only the local connectivity but also the higher-order topology of the structural connectome. Specifically, our findings suggest that cytoarchitecture may constrain the traversal of structural pathways to engender efficient bottom-up signal routing over the hierarchy. Thus, it appears that cytoarchitecture not only predicts which pairs of regions are connected (i.e., “like connects with like” cf. the *structural model*^1–4^), but also the spatial embedding of senders and receivers in the brain^50,67,68^.

### Energy asymmetries link to changes in intrinsic neuronal timescales and effective connectivity

Recent work has shown that the spatial patterning of regions’ intrinsic neuronal timescales correlate with the patterning of the T1w/T2w ratio^10^, suggesting that the brain’s timescale hierarchy reflects its cytoarchitectonic hierarchy. Here, we found that the asymmetries in trans-hierarchical state transitions were tightly coupled to differences in state-level intrinsic neuronal timescales. Specifically, the easier a bottom-up state was to complete (compared to its top-down counterpart) the more the timescale of the target state lengthened compared to the initial state. Lengthening timescales are thought to be associated with progressive changes to longer temporal receptive windows, which in turn is thought to underpin shifts from segregated to integrated functional processing^69^. Thus, our findings show that the topology of the structural connectome may be wired to support the progressive integration of lower-order properties of our environment into higher-order percepts and cognitions. Our findings also serve as a functional validation of our network control model; we observed a positive correlation between energy asymmetries and asymmetries in effective connectivity, which is consistent with past literature^50^. Thus, our findings contribute to a growing body of evidence demonstrating that asymmetric signal routing is measurable from the topology of the connectome, despite being derived from an undirected description of brain connectivity.

### Energy asymmetries refine systematically throughout youth

The effects of development on connectome topology are increasingly well studied^42,70–72^, including with network control theory where the amount of control energy required to activate the executive function system (from baseline) has been shown to decrease throughout youth^73^. This observation is consistent with the current study, wherein the energy associated with trans-hierarchical state transitions also reduced4 throughout youth. Here, we provide a key extension to prior work that deepens our understanding of these developmental energy effects; we observed that age effects were (i) stronger for top-down compared to bottom-up energy and (ii) concentrated in state transitions with the most pronounced energy asymmetries. These findings suggest that maturation throughout youth alters the balance between bottom-up and top-down signal propagation, refining the connectome towards an equilibrium between the two. This interpretation is consistent with a staging account of neurodevelopment that suggests that lower-order connections are refined earlier in development compared to their higher-order counterparts^3,39–41^. That is, the energy asymmetry we observed might reflect the relatively advanced refinement of lower-order connections that is already well underway by 8 years of age (the youngest in our sample). In turn, the stronger age effect observed for top-down energy might reflect the relatively delayed onset of refinement of higher-order connections that may be occurring within the age range of our sample. Examining how our results present on either side of the age range of the PNC, as well as whether it is supported by longitudinal data, will be a critical avenue for future research.

## Limitations

Similar to our recent work^74^, a limitation of this study is the use of a linear model of neuronal dynamics to estimate signal propagation across the S-F axis. While this assumption is an over-simplification of brain dynamics, linear models explain variance in the slow fluctuations in brain activity recorded by fMRI^75,76^, suggesting that they successfully approximate the kinds of data commonly used to examine brain function. An additional limitation is the use of a single map of cytoarchitecture to define brain states, which precluded us from defining participant-specific states. As mentioned above, previous work has shown that the T1w/T2w ratio forms a reasonable proxy of the S-F axis of cytoarchitecture that is measurable *in vivo*^5^. However, while the PNC includes T1-weighted imaging, it does not include T2-weighted imaging^53^, which prevented us from estimating the T1w/T2w ratio in our sample. Replication of our findings using participants’ T1w/T2w maps is warranted given well-known individual variability in the spatial patterning of cortical structural features. However, this approach must be weighed against the fact the T1w/T2w ratio is imperfectly correlated with cytoarchitecture. Thus, such replication efforts must consider the trade-off between the value of individual variability and the cost of potentially disconnecting from the relevant underlying neurobiology (i.e., cytoarchitecture).

## Conclusions

Our results demonstrate that cytoarchitecture may constrain network topology in such a way as to induce asymmetries in signal propagation across the cortical hierarchy. Specifically, we found that bottom-up trans-hierarchical state transitions were easier to complete than their top-down counterparts, that asymmetries correlated with changes to neuronal timescales, that control signals tracked the sensory-fugal axis, and that asymmetries reduced with age in youth. Collectively, our work highlights that variation in the properties of cortical microstructure that govern extrinsic connectivity may guide the formation of macroscopic connectome topology.

## Materials and Methods

### Participants

Participants included 793 individuals from the Philadelphia Neurodevelopmental Cohort^77,78^, a community-based study of brain development in youths aged 8 to 22 years^79,80^. The institutional review boards of both the University of Pennsylvania and the Children’s Hospital of Philadelphia approved all study procedures. The neuroimaging sample of the PNC consists of 1,601 participants^77^. From this original sample, 156 were excluded due to the presence of gross radiological abnormalities distorting brain anatomy or due to a medical history that might impact brain function. Next, a further 159 participants were excluded because they were taking psychoactive medication at the time of study. An additional 466 individuals were excluded because they did not pass rigorous manual and automated quality assurance for their T1-weighted scan^81^, their diffusion scan^82^, or their resting-state functional magnetic resonance imaging (rs-fMRI) scan^83,84^. Finally, 27 participants were excluded owing to the presence of disconnected regions in their structural connectivity matrix (see section entitled *Structural connectome construction* below). This process left a final sample of 793 participants.

### Imaging data acquisition

MRI data were acquired on a 3 Tesla Siemens Tim Trio scanner with a 32-channel head coil at the Hospital of the University of Pennsylvania. Diffusion weighted imaging (DWI) scans were acquired via a twice-refocused spin-echo (TRSE) single-shot echo-planar imaging (EPI) sequence (TR=8100 ms, TE=82 ms, FOV=240mm^2^/240mm^2^; Matrix=RL: 128, AP: 128, Slices: 70, in-plane resolution of 1.875 mm^2^; slice thickness=2 mm, gap=0; flip angle=90°/180°/180°, 71 volumes, GRAPPA factor=3, bandwidth=2170 Hz/pixel, PE direction=AP). The sequence utilized a four-lobed diffusion encoding gradient scheme combined with a 90-180-180 spin-echo sequence designed to minimize eddy-current artifacts^53^. The sequence consisted of 64 diffusion-weighted directions with *b*=1000 s/mm^2^ and 7 interspersed scans where *b*=0 s/mm^2^. The imaging volume was prescribed in axial orientation and covered the entire brain.

In addition to the DWI scan, a B0 map of the main magnetic field was derived from a double-echo, gradient-recalled echo (GRE) sequence, allowing for the estimation and correction of field distortions. Prior to DWI acquisition, a 5-min magnetization-prepared, rapid acquisition gradient-echo T1-weighted (MPRAGE) image (TR=1810 ms, TE=3.51 ms, FOV=180 × 240 mm, matrix 256 × 192, effective voxel resolution of 0.94 × 0.94 × 1 mm) was acquired for each participant.

Finally, approximately 6 minutes of rs-fMRI data was acquired using a blood oxygen level-dependent (BOLD-weighted) sequence (TR=3000 ms; TE=32 ms; FoV=192 × 192 mm; resolution 3 mm isotropic; 124 volumes). These data were used primarily to generate the principal cortical gradient of functional connectivity discussed in the main text^12^ (see section entitled *Principal gradient of functional connectivity* below).

### Imaging data quality control

All DWI and T1-weighted images underwent rigorous quality control by highly trained image analysts (see Roalf et al. (2016) and Rosen et al. (2018) for details on DTI and T1-weighted imaging, respectively). Regarding the DWI acquisition, all 71 volumes were visually inspected and evaluated for the presence of artifacts. Every volume with an artifact was marked as contaminated and the fraction of contaminated volumes was taken as an index of scan quality. Scans were marked as ‘poor’ if more than 20% of volumes were contaminated, ‘good’ if more than 0% but less than 20% of volumes were contaminated, and ‘great’ if 0% of volumes were contaminated. Regarding the T1-weighted acquisition, images with gross artifacts were considered ‘unusable’; images with some artifacts were flagged as ‘usable’; and images free of artifact were marked as ‘superior’. As mentioned above in the section entitled *Participants*, 466 individuals were removed due to quality. Of these, 318 individuals were removed due to either ‘poor’ DWIs or ‘unusable’ T1-weighted images. In the final sample of 793 participants, a total of 535 participants had diffusion tensor images identified as ‘great’, with the remaining identified as ‘good’, and 701 participants had T1-weighted images identified as ‘superior’, with the remaining identified as ‘usable’. Regarding the rs-fMRI data, as in prior work^83,84^, the remaining 148 of 466 excluded participants were removed either because their mean relative root mean square (RMS) framewise displacement was higher than 0.2 mm or their scan included more than 20 frames with motion exceeding 0.25 mm.

### Structural image processing

Structural image processing was carried out using tools included in ANTs^85^. The *buildtemplateparallel* pipeline from ANTs^86^ was used to create a study-specific T1-weighted structural template with 120 participants that were balanced on sex, race, and age. Structural images were processed in participants’ native space using the following procedure: brain extraction, N4 bias field correction^87^, Atropos tissue segmentation^88^, and SyN diffeomorphic registration^86,89^.

### Diffusion image processing

For each participant, a binary mask was created by registering the standard fractional anisotropy mask provided by FSL (FMRIB58 FA) to the participant’s mean *b*=0 reference image using FLIRT^90^. To correct for eddy currents and head motion, this mask and the participant’s diffusion acquisition was passed to FSL’s *eddy*^91^ (version 5.0.5). Diffusion gradient vectors were subsequently rotated to adjust for the motion estimated by *eddy*. Distortion correction was conducted via FSL’s FUGUE^92^ using the participant’s field map, estimated from the B0 map.

### rs-fMRI processing

State-of-the-art processing of functional data is critical for valid inference^93^. Thus, functional images were processed using a top-performing preprocessing pipeline implemented using the eXtensible Connectivity Pipeline (XCP) Engine^83^, which includes tools from FSL^92,94^ and AFNI^95^. This pipeline included (1) correction for distortions induced by magnetic field inhomogeneity using FSL’s FUGUE utility, (2) removal of 4 initial volumes, (3) realignment of all volumes to a selected reference volume using FSL’s MCFLIRT, (4) interpolation of intensity outliers in each voxel’s time series using AFNI’s 3dDespike utility, (5) demeaning and removal of any linear or quadratic trends, and (6) co-registration of functional data to the high-resolution structural image using boundary-based registration. Images were de-noised using a 36-parameter confound regression model that has been shown to minimize associations with motion artifact while retaining signals of interest in distinct sub-networks^83,96^. This model included the six framewise estimates of motion, the mean signal extracted from eroded white matter and cerebrospinal fluid compartments, the mean signal extracted from the entire brain, the derivatives of each of these nine parameters, and quadratic terms of each of the nine parameters and their derivatives. Both the BOLD-weighted time series and the artifactual model time series were temporally filtered using a first-order Butterworth filter with a passband of 0.01–0.08 Hz^97^.

### Imaging-derived nuisance covariates

In our analyses of individual differences, we used total brain volume and mean in-scanner motion as imaging-derived nuisance covariates. Total brain volume was generated from the T1-weighted images using ANTs. In-scanner head motion was estimated for each participant from their DWI sequence as relative framewise displacement^82^. Specifically, rigid-body motion correction was applied to the seven high quality *b*=0 images interspersed throughout the diffusion acquisition. Once estimated, framewise displacement was averaged across time to create a single measure for each participant.

### Structural connectome construction

For each participant, whole-brain deterministic fiber tracking was conducted using DSI Studio^98^ with a modified fiber assessment by continuous tracking (FACT) algorithm with Euler interpolation. A total of 1,000,000 streamlines were generated for each participant that were between 10mm and 400mm long. Fiber tracking was performed with an angular threshold of 45° and step size of 0.9375 mm. Next, following our previous work^74^, the number of streamlines intersecting region *i* and region *j* in a 200-parcel cortical parcellation^99^ was used to weight the edges of an undirected adjacency matrix, ***A*** (see **Fig. S5** for sensitivity analyses covering different parcellation resolutions and definitions). Note that ***A***_*ij*_ = 0 for *i* = *j*. This process yielded 793 subject-specific ***A*** matrices that were used in subject-level analyses reported in the main text (i.e., **Fig. 6**). Our primary analyses, however, were based on a group-averaged ***A*** matrix. To obtain this ***A*** matrix, we averaged over the entries of the individuals’ ***A*** matrices and thresholded using an edge consistency-based approach^100^. Specifically, edges in the group-averaged ***A*** matrix were only retained if non-zero edge weights were present in at least 60% of participants’ ***A*** matrices^101^. If not, edges were set to zero. This process yielded a group-averaged structural connectome with a sparsity value of approximately 8%. This group-averaged structural connectome was used for analyses reported in **Figs. 2, 3, 4**, and **5**. See **Fig. S6** for sensitivity analyses spanning a range of consistency thresholds and corresponding sparsity values.

## Trans-hierarchical state transitions

### Cortical hierarchies

Below we describe two views of the cortical hierarchy that were used in the present study to examine trans-hierarchical state transitions.

### Sensory-fugal axis of cytoarchitecture

We primarily characterized the cortical hierarchy using the gradient of cytoarchitectonic similarity developed in previous work^5,66^ and disseminated as part of the BigBrainWarp toolbox^6^. Specifically, from BigBrainWarp, we retrieved the histological gradient (‘Hist-G2’) corresponding to the sensory-fugal (S-F) axis of cytoarchitecture stored in *fsaverage* space. Next, we averaged over the vertex values within each of our 200 cortical parcels (see section entitled *Structural connectome construction* above). This process resulted in a 200 × 1 vector describing regions’ positions along the S-F axis of cytoarchitectonic similarity.

### Principal gradient of functional connectivity

As stated above and in the main text, our primary constituent of the cortical hierarchy was the S-F axis of cytoarchitectonic similarity. To test the specificity of our primary results, we also examined another view of the cortical hierarchy: the principal gradient of functional connectivity^12^. This gradient situates unimodal sensorimotor cortex at one end and transmodal association cortex at the other. Conceptually, this approach amounts to a dimensionality reduction technique that positions regions with similar functional connectivity profiles near to one another, and positions regions with dissimilar functional connectivity profiles distant from one another. Here, as in our previous work^74^, we generated this gradient using whole-brain resting-state functional connectivity obtained from the PNC data (see section entitled *rs-fMRI processing* above). Specifically, for each participant, processed rs-fMRI timeseries were averaged regionally and a Pearson correlation coefficient was estimated between each pair of regional timeseries to generate a functional connectome. Correlation coefficients were normalized using Fisher’s *r*-to-*z* transform, and then connectomes were averaged over participants. The principal gradient of functional connectivity was generated from this group-average functional connectome using a diffusion map embedding implemented in the *BrainSpace* toolbox^102^. We selected the first gradient output from this approach, which was closely aligned to that observed previously^12^. Note that this gradient is the same as that reported in our previous work^74^. This process resulted in a 200 × 1 vector that describes regions’ positions along the unimodal-to-transmodal (U-T) axis of functional connectivity.

### Hierarchical brain states

As discussed in the main text and illustrated in **Fig. 1**, we divided our 200 × 1 S-F axis of cytoarchitecture—as well as the U-T axis of functional connectivity—into 20 evenly sized (*n*=10) and non-overlapping sets of brain regions that traversed up the cortical hierarchy. This procedure yielded 20 groups of cortical regions that differed based on their position along the S-F (U-T) axis. Thus, regions within each group had similar profiles of cytoarchitecture (functional connectivity) while regions between groups had dissimilar profiles of cytoarchitecture (functional connectivity). Moreover, this dissimilarity increased with greater distance between pairs of groups along the S-F (U-T) axis. These 20 groups of regions formed the brain states that we used in the network control theory analysis (see section entitled *Network control theory* below), thus allowing us to model transitions between states moving up and down the cortical hierarchy. See **Fig. S7** for sensitivity analyses covering different set sizes for brain states.

### Network control theory

To model trans-hierarchical state transitions, we employed tools from network control theory^51,52,62,63^. Given an ***A*** matrix as input (either group-averaged or individual; see section entitled *Structural connectome construction* above), we first apply the following normalization:

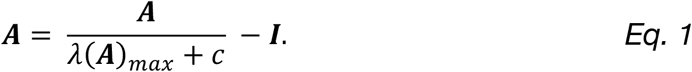

Here, λ(***A***)_*max*_ is the largest eigenvalue of ***A***, *c* = 1 to ensure system stability, and ***I*** denotes the identity matrix of size *N* × *N*. In our analyses, *N* is equal to the number of brain regions, which is 200. Within this normalized ***A*** matrix, we allow each node of the network to carry a real value representing that node’s activity. These values are represented in *x* and collectively describe the pattern of whole-brain activity as it changes over time. Next, we use a simplified noise-free linear continuous-time and time-invariant model of network dynamics:

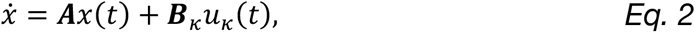

where *x*(*t*) is a *N* × 1 vector that represents the state of the system at time *t*. The matrix ***B***, identifies the control input weights, which by default we set to the *N* × *N* identity matrix to compute unweighted energy (see section entitled *Minimizing transition energy through optimized control weights* below for the weighted case).

Given this model of the dynamics, we compute the control inputs, *u*,(*t*), that drive the system from some initial state, *x*_0_, to some target state, *x*^*^, in a finite amount of time *T* = 1. Here, initial and target states were constructed using the 20 non-overlapping groups of 10 brain regions spanning the S-F axis (see above section entitled *Hierarchical brain states*). That is, each initial or target state was defined as an *N* × 1 vector within which 10 elements that represented cytoarchitecturally similar areas contained a 1, and the remaining elements contained a 0. Among the many possible inputs, we chose the *minimum energy*^51,103^ input which minimizes a quadratic cost on the inputs, such that

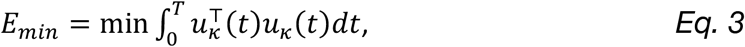

subject to *Eq. 2*. To compute the minimum energy, we construct a useful mathematical object called the *controllability Gramian*, given by

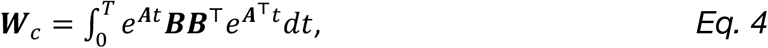

where *e*^***A****t*^ is the time-dependent matrix exponential of the matrix ***A***, and is also the *impulse response* of the system that governs the natural evolution of system dynamics. Then, the minimum energy is given by

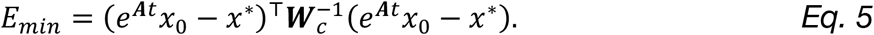

Intuitively, the quantity in the parentheses measures the difference between the natural evolution of the system from the initial condition, *e*^***A****t*^*x*_0_, and the target state, *x*^*^. This difference is precisely the difference for which the control input *u*,(*t*) needs to compensate, and the projection of this difference onto 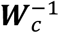 yields the minimum energy for providing such compensation^52^.

### Transition energy and energy asymmetries

We used the above derivation of minimum control energy to compute a *k* × *k* transition energy matrix, *T*_*E*_. Elements of *T*_*E*_ quantified the energy (*E*) required to transition between all possible pairs of *k* = 20 brain states, where brain states were based on the subsets of regions sampled along the S-F axis of cytoarchitecture outlined above (see section entitled *Cortical hierarchies*). As mentioned in the main text and above, the hierarchically ordered nature of our brain states endowed *T*_*E*_ with a distinction between transitions moving up the hierarchy (bottom-up energy) from those moving down the hierarchy (top-down energy). Further, these bottom-up and top-down transition energies were naturally compartmentalized into the upper and lower triangles of *T*_*E*_, respectively. Hence, asymmetries between bottom-up and top-down energy for all state pairs were calculated as 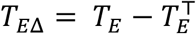. Note that unlike *T*_*E*_, *T*_*E*Δ_ is symmetrical; thus, only the upper triangle was carried forward for asymmetry analysis.

### Null network models

In **Fig. 2B** in the main text, we showed that *T*_*E*Δ_ was negatively correlated with the distance that separated states along the hierarchy. We compared this empirically observed correlation with hierarchy distance to those expected under two spatially embedded null models^54^. Alongside preserving the spatial embedding of network nodes, these null models randomly rewired the network while preserving either the edge distribution or the strength distribution of the network. For each of these null models, we produced 5,000 rewired networks derived from the group-averaged structural connectome (see section entitled *Structural connectome construction* above) using publicly available code (https://github.com/breakspear/geomsurr). Then, to generate an empirical null distribution, upon each rewired network we recomputed *T*_*E*_, *T*_*E*Δ_, and the corresponding hierarchy distance correlation with *T*_*E*Δ_. Finally, *p*-values were estimated as the probability that the magnitude of the observed distance correlation occurred under a given null.

### Uncontrolled dynamics

In addition to examining the energy required to complete state transitions between specific state-pairs, we also examined how uncontrolled dynamics spread naturally across the cortex from each of our cytoarchitectonic brain states. Specifically, for each brain state, we set the constituent regions’ activity to 1 and all other regions’ activity to

0. Then, we allowed the activity to diffuse in an uncontrolled manner along the networks’ edges over time according to 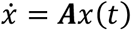; this approach stands in contrast to the approach that we have discussed thus far of forcing activity to flow from one state to another via a set of control signals. As mentioned in the main text, we analyzed these natural dynamics in two ways. First, for each seed brain state and time point, *t*, we correlated the pattern of simulated activity at each node with the sensory-fugal axis of cytoarchitecture. Note that this correlation was computed excluding the regions that made up a given seed state (i.e., where activity was propagating from). Thus, correlations were not driven by activity leaving a given brain state, but rather only reflected where in the brain that activity flowed to. Results of this analysis are shown in **Fig. 3B** and **3C**.

Second, we averaged activity over regions within each brain state, creating a 20 × *n* matrix of state-averaged activity over time (see **Fig. 3D** for examples). We repeated this process using each brain state to seed activity to create a 20 × 20 × *n* activity matrix, where the first dimension denoted the brain state from which activity was seeded and where dimensions two and three stored how activity spread to all other brain states over time (including the seed state itself). Next, for each seed state, we estimated the correlation between the position along the S-F axis of the remaining brain states and the point in time when activity peaked within those states. Thus, a positive correlation indicated that activity propagated up the hierarchy from a given seed state while a negative correlation indicated that activity propagated down the hierarchy (see **Fig. 3E**). Lastly, we compared the magnitude of these correlations between pairs of seed brain states using 1,000 bootstrapped versions of our group-averaged structural connectome. Specifically, we reproduced our group-averaged structural connectome for each of 1,000 bootstrapped samples of our 793 participant connectomes. Then, using these 1,000 bootstrapped connectomes, we re-simulated the spread of uncontrolled dynamics and re-estimated the propagation correlations. This procedure allowed us to estimate 95% confidence intervals for the magnitude of the observed propagation correlations, which in turn allowed us to compare their size (see **Fig. 3F**).

### Intrinsic neuronal timescales

As mentioned in the main text, we sought to validate our transition energy analysis in functional data using *intrinsic neuronal timescales* derived from electrocorticography (ECoG) data. Thus, we compared energy asymmetries from our NCT analysis with differences between brain states’ intrinsic timescales. Following previous work^10^, we estimated regions’ intrinsic timescales using the time constant (τ) of an exponential decay function fitted to the autocorrelation function of ECoG timeseries. Specifically, we downloaded sensor-level τ data processed by Gao *et al*. (https://github.com/rdgao/field-echos/data/df_human.csv) and, using the provided MNI coordinates, matched each sensor to our parcellation (200 Schaefer parcels); matching was done by finding the smallest Euclidean distance between each sensor and the centroid of each parcel. We then averaged τ over sensors within each parcel as well as over regions within each cytoarchitectonic brain state. This process generated state-level τ values that were then subtracted to produce τ_Δ_, a matrix of change in τ between all pairs of brain states.

### Effective connectivity

As mentioned in the main text, we sought to validate our transition energy analysis in functional data using *effective connectivity* derived from rs-fMRI data. Thus, we compared energy asymmetries from our NCT analysis with asymmetries in effective connectivity. As per our previous work^104^, effective connectivity was estimated using a spectral version of dynamic causal modeling (spDCM)^60,61^ implemented in SPM12 r7765 (Wellcome Trust Centre for Neuroimaging, London, UK). To generate timeseries for modeling effectivity connectivity, we first averaged participants’ processed rs-fMRI data across the regions that comprised each cytoarchitectonic brain state. This process yielded one timeseries of 120 volumes per subject per brain state. Next, owing to the low number of volumes in our rs-fMRI acquisition, we deployed an averaging and concatenation approach that yielded a single group-averaged timeseries of 1200 volumes for each brain state. This process proceeded as follows. First, we randomly excluded 3 participants from our sample to retain 790 participants. Second, we divided our sample of 790 participants into 10 equally sized groups (*n*=79) and averaged the state-level rs-fMRI timeseries across participants within each group separately. Finally, we concatenated these group-averaged timeseries end-to-end across the 10 groups. This process yielded resting-state timeseries for each brain state with 1200 volumes that represented averages over distinct subsets of participants taken from our sample. These timeseries were used as inputs to the spDCM algorithm, together with a fully connected model of coupling strengths, enabling the estimation of effective connectivity between all cytoarchitectonic brain states spanning the S-F axis. As per our primary analysis of transition energies, effective connectivity estimates were trivially grouped into bottom-up and top-down, and were then subtracted to create an effective connectivity asymmetry matrix.

### Minimizing transition energy through optimized control weights

Our primary analyses involved examining uniformly weighted transition energies, where all nodes of the dynamical system were assigned control weights equal to 1 (i.e., setting the diagonal entries of ***B***, in *Eq. 2* to the *N* × *N* identity matrix). This uniform weighting meant that all brain regions were endowed with the same degree of control over all *k* × *k* state transitions. However, as discussed in the main text (see section entitled *Optimized control weights increase energy asymmetries and track the sensory-fugal axis of cytoarchitecture*), we were also interested in examining regional variation in facilitating trans-hierarchical state transitions.

To achieve this goal, we systematically perturbed each region’s degree of control over the system and measured the corresponding change in transition energies. Specifically, for each brain region, *i*, we recomputed *T*_*E*_ after adding a constant amount of additional control to the corresponding diagonal element of ***B***, (the remaining diagonal entries were left equal to 1). This process generated a *k* × *k* × 200 matrix of perturbed transition energies, *P*_*E*_. Next, for each perturbed region (dimension 3 of *P*_*E*_), we subtracted the perturbed transition energies from the uniformly weighted energies (*T*_*E*_) to create *P*_*E*Δ_, a *k* × *k* × 200 matrix of perturbed transition energy deltas. For each state transition, this subtraction yielded a 200 × 1 vector that quantified how perturbing each node of the system one at a time—by a constant arbitrary amount—impacted transition energy. Note that increasing the influence of a single node’s control necessarily reduces energy; the task of completing a state transition is easier for the model when any node in ***B***, is granted a greater degree of control over the system, leading to lower energy. Accordingly, all values in *P*_*E*Δ_ were positive and the magnitude of these deltas encoded the relative importance of each region to completing a specific state transition, with regions with larger deltas being more important.

To assess correspondence with the S-F axis, we calculated the Spearman rank correlation between perturbed deltas for each state transition and the gradient of cytoarchitecture (see **Fig. S3**). Next, we re-estimated *T*_*E*_ (and *T*_*E*Δ_) one more time using each state transition’s vector of perturbed deltas as optimized control weights. This process yielded optimized trans-hierarchical transition energies and optimized energy asymmetries. Finally, to assess whether the size of the mean *T*_*E*Δ_ was significantly different for optimized weights compared to uniform weights, we derived *T*_*E*Δ_ for both weight sets using bootstrapped group-averaged connectomes (see section entitled *Uncontrolled dynamics* above) and assigned 95% confidence intervals to the mean *T*_*E*Δ_.

### Age effects

As mentioned in the main text, we sought to link subject-specific energy asymmetries with age to examine developmental effects. To achieve this goal, we derived *T*_*E*_ (and *T*_*E*Δ_) from each participant’s ***A*** matrix (see section entitled *Structural connectome construction* above) using optimized control weights. Note that the process of computing optimized transition energies was performed on a subject-specific basis using subject-specific optimized control weights; this was done by applying the above perturbation procedure (see section entitled *Minimizing transition energy through optimized control weights*) to each participant’s ***A*** matrix separately (see **Fig. S8** for correlations between subject-specific optimized weights and the gradient of cytoarchitecture). Next, for each state transition, we calculated the Pearson’s correlation between *T*_*E*_ and age, while controlling for sex, total brain volume, and in-scanner motion (see section entitled *Imaging-derived nuisance covariates* above). We repeated this process for average bottom-up and top-down energy, where energy was averaged over the upper and lower triangles of each participant’s *T*_*E*_ matrix, respectively.

In addition to estimating within-sample age effects, we also sought to test whether energy asymmetries could be used to predict participants’ ages in out-of-sample testing. To achieve this, we assembled the upper triangle of each subject’s *T*_*E*Δ_ matrix into a 793 × 190 feature table, *X*. To ensure normality, columns of *X* were normalized using an inverse normal transformation^105,106^. Then, we used a cross-validated ridge regression model implemented in *scikit-learn*^107^ with default parameters (*α* = 1) to predict participants’ ages (*y*). Specifically, we assessed out-of-sample prediction performance using 10-fold cross-validation scored by root mean squared error (RMSE) and by the correlation between the true *y* and predicted *y*. Note, as per *scikit-learn* defaults, to standardize the interpretation of both scoring metrics as higher scores represent better performance, we flipped the sign for RMSE and examined negative RMSE.

Models were trained using all columns of *X* as input features and scoring metrics were each averaged across folds. As above, we included sex, total brain volume, and in-scanner motion as nuisance covariates. Nuisance covariates were controlled for by regressing their effect out of *X* before predicting *y*. Within each fold, nuisance covariates were fit to the training data and applied to the test data to prevent leakage. Subsequently, we applied principal component analysis (PCA) to reduce the dimensionality of *X*, retaining enough PCs to explain 80% of the variance in the data. Finally, owing to evidence that prediction performance can be biased by the arbitrariness of a single split of the data^108^, we repeated 10-fold cross-validation 100 times, each time with a different random 10-fold split. This process yielded a distribution of 100 mean negative RMSE values and 100 mean correlations between true *y* and predicted *y*.

Our above prediction model generated robust estimates of prediction performance, but it did not examine whether prediction performance was itself significant. To test whether prediction performance was better than chance, we compared point estimates of each of our scoring metrics—taken as the mean over the 100 values—to the distribution of values obtained from permuted data. Specifically, we subjected the point estimates of our scoring metrics to 5,000 random permutations, wherein the rows (i.e., participants) of *y* were randomly shuffled. The associated *p*-values were assigned as the proportion of permuted scores that were greater than or equal to our true scores.

## Citation diversity statement

Recent work in several fields of science has identified a bias in citation practices such that papers from women and other minority scholars are under-cited relative to the number of such papers in the field^109–117^. Here we sought to proactively consider choosing references that reflect the diversity of the field in thought, form of contribution, gender, race, ethnicity, and other factors. First, we obtained the predicted gender of the first and last author of each reference by using databases that store the probability of a first name being carried by a woman^113,118^. By this measure (and excluding self-citations to the first and last authors of our current paper), our references contain 7.82% woman(first)/woman(last), 12.25% man/woman, 16.98% woman/man, and 62.95% man/man. This method is limited in that a) names, pronouns, and social media profiles used to construct the databases may not, in every case, be indicative of gender identity and b) it cannot account for intersex, non-binary, or transgender people. Second, we obtained predicted racial/ethnic category of the first and last author of each reference by databases that store the probability of a first and last name being carried by an author of color^119,120^. By this measure (and excluding self-citations), our references contain 6.03% author of color (first)/author of color(last), 19.77% white author/author of color, 20.93% author of color/white author, and 53.27% white author/white author. This method is limited in that a) names and Florida Voter Data to make the predictions may not be indicative of racial/ethnic identity, and b) it cannot account for Indigenous and mixed-race authors, or those who may face differential biases due to the ambiguous racialization or ethnicization of their names. We look forward to future work that could help us to better understand how to support equitable practices in science.

## Acknowledgments

LP was supported by the National Institute Of Mental Health of the National Institutes of Health under Award Number K99MH127296 and a 2020 NARSAD Young Investigator Grant from the Brain & Behavior Research Foundation. The content is solely the responsibility of the authors and does not necessarily represent the official views of the National Institutes of Health. This study was additionally supported by the National Institute of Mental Health (Grant Nos. R21MH106799 [to DSB and TDS], R01MH113550 [to TDS and DSB], and RF1MH116920 [to TDS and DSB]) and the Swartz Foundation. Additional support was provided by the John D. and Catherine T. MacArthur Foundation (to DSB) as well as grant Nos. R01MH120482 (to TDS), R01MH107703 (to TDS), R01MH112847 (to TDS and RTS), R01MH120482 (to TDS), R37MH125829 (to TDS), R01EB022573 (to TDS), R01MH107235 (to RCG), R01 MH119219 (to RCG and REG), R01MH119185 (to DRR), R01MH120174 (to DRR), R01MH113565 (to DHW), and the Penn-CHOP Lifespan Brain Institute. JZK was supported by the National Science Foundation DGE-1321851. The PNC was supported by RC2MH089983 and RC2MH089924.

## Author contributions

Conceptualization: L.P., D.S.B, and T.D.S.; Methodology: L.P., J.Z.K, J.S., R.T.S., and

D.S.B.; Software: L.P., J.Z.K, J.S., and M.O.; Formal analysis: L.P.; Data curation: M.E.C., M.C., R.E.G, R.C.G., T.M.M., D.R.R., R.T.S., D.H.W, and T.D.S; Writing—original draft: L.P.; Writing—reviewing and editing: L.P., J.Z.K, J.S., M.E.C., M.C., R.E.G, R.C.G., T.M.M., M.O., D.R.R., R.T.S., D.H.W, T.D.S, and D.S.B.; Visualization: L.P. and M.O.

## Code availability

The PNC data are publicly available in the Database of Genotypes and Phenotypes: accession number: phs00607.v3.p2; https://www.ncbi.nlm.nih.gov/projects/gap/cgi-bin/study.cgi?study_id=phs000607.v3.p2. All analysis code is available at https://github.com/lindenmp/nct_hierarchy.

## Supplementary Materials for

**Figure S1.**
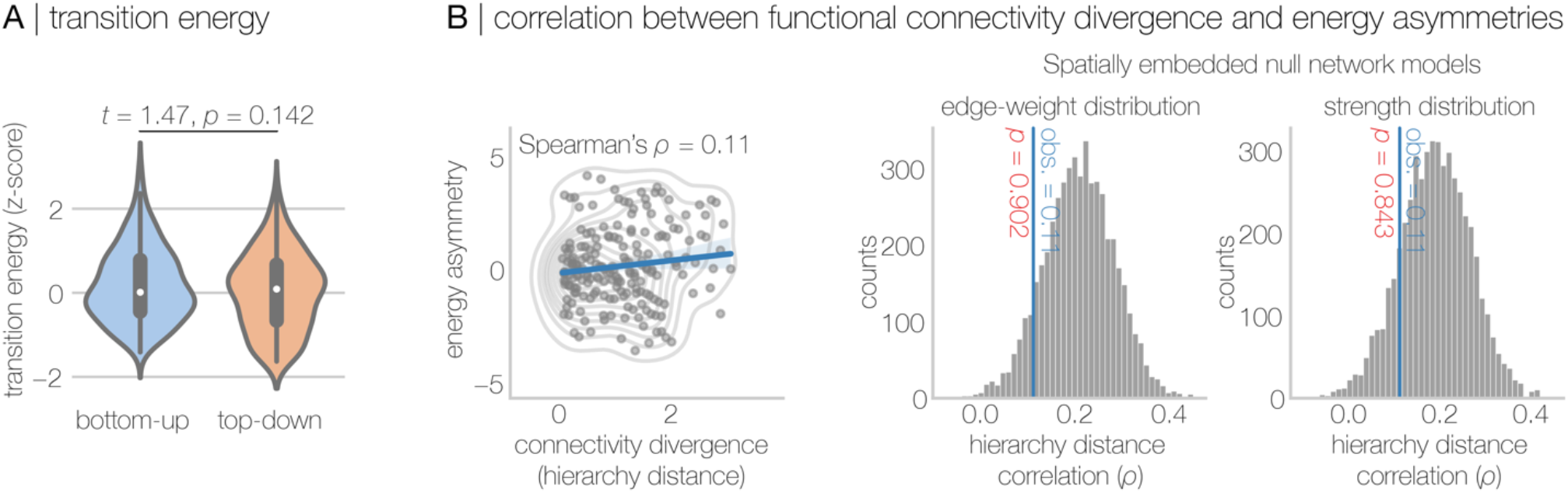
Counterpart to Figure 2 using the principal gradient of functional connectivity (Margulies et al., 2016) to define brain states instead of the sensory-fugal axis of cytoarchitecture (Paquola et al., 2019). **A**, No significant differences between bottom-up and top-down energy were observed when the principal gradient of functional connectivity was used to define brain states. **B**, The distance along the functional connectivity gradient separating initial and target states was not significantly correlated with energy asymmetry.

**Figure S2.**
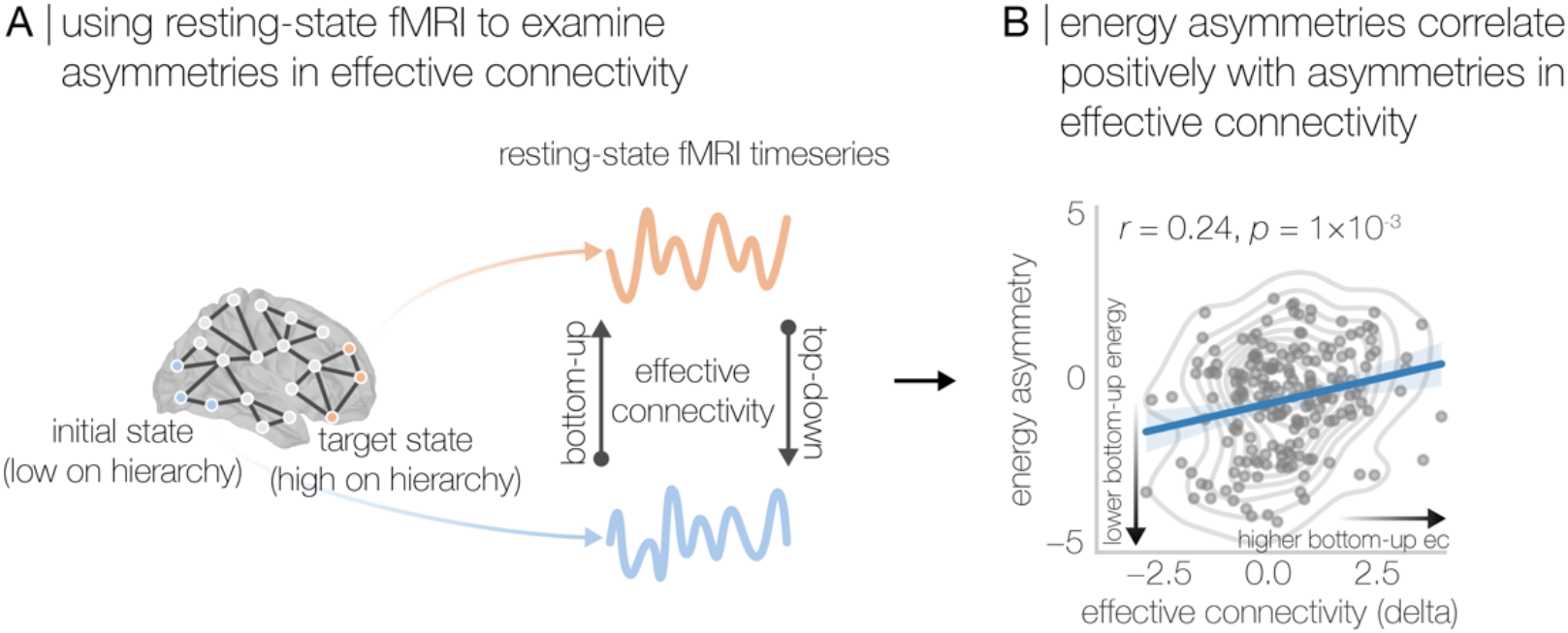
Energy asymmetries correlate with asymmetries in effective connectivity measured from resting-state functional magnetic resonance imaging. **A**, We used resting-state functional magnetic resonance imaging to examine asymmetries in effective connectivity (estimated using dynamic causal modeling) between our cytoarchitectonic brain states (see Methods). **B**, Energy asymmetries between brain states correlated positively with asymmetries in effective connectivity between brain states. This result shows that for state transitions where bottom-up transition energy was lower than top-down (negative energy asymmetry) the same was true for effective connectivity and *vice versa*.

**Figure S3.**
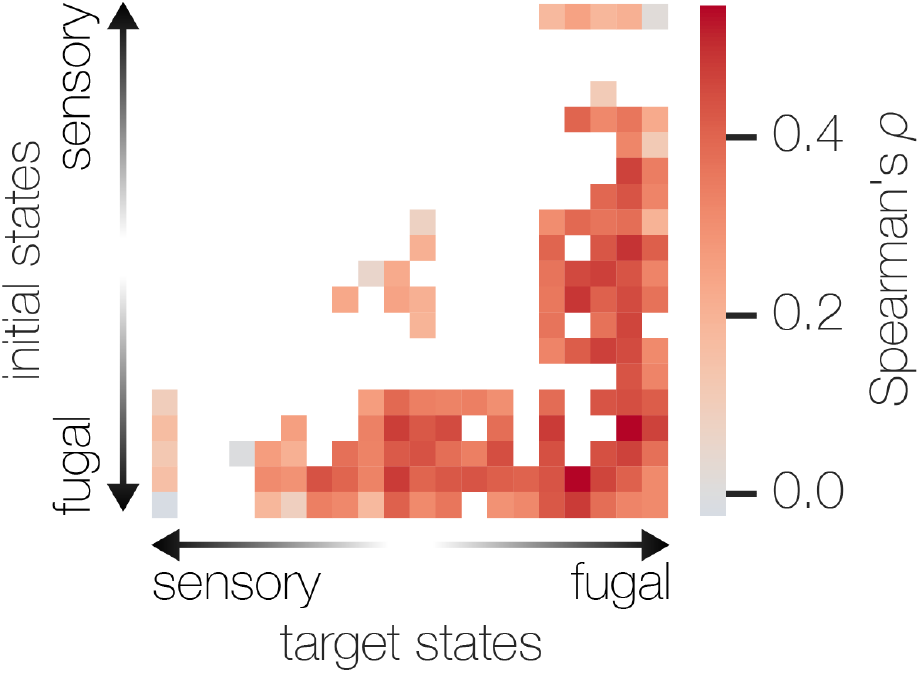
Significant correlations between the sensory-fugal axis of cytoarchitecture and optimized control weights for each state transition. Optimized control weights correlated positively with the cortical hierarchy of cytoarchitecture, indicating that regions higher on the hierarchy were more important for control. The *p*-values for correlations were estimated using a null network model that preserved the spatial embedding of brain region as well as the strength distribution. The *p*-values were corrected for multiple comparisons using the Benjamini-Hochberg False Discovery Rate (Benjamini & Hochberg, 1995). Significance was determined as *p*_FDR_<0.05.

**Figure S4.**
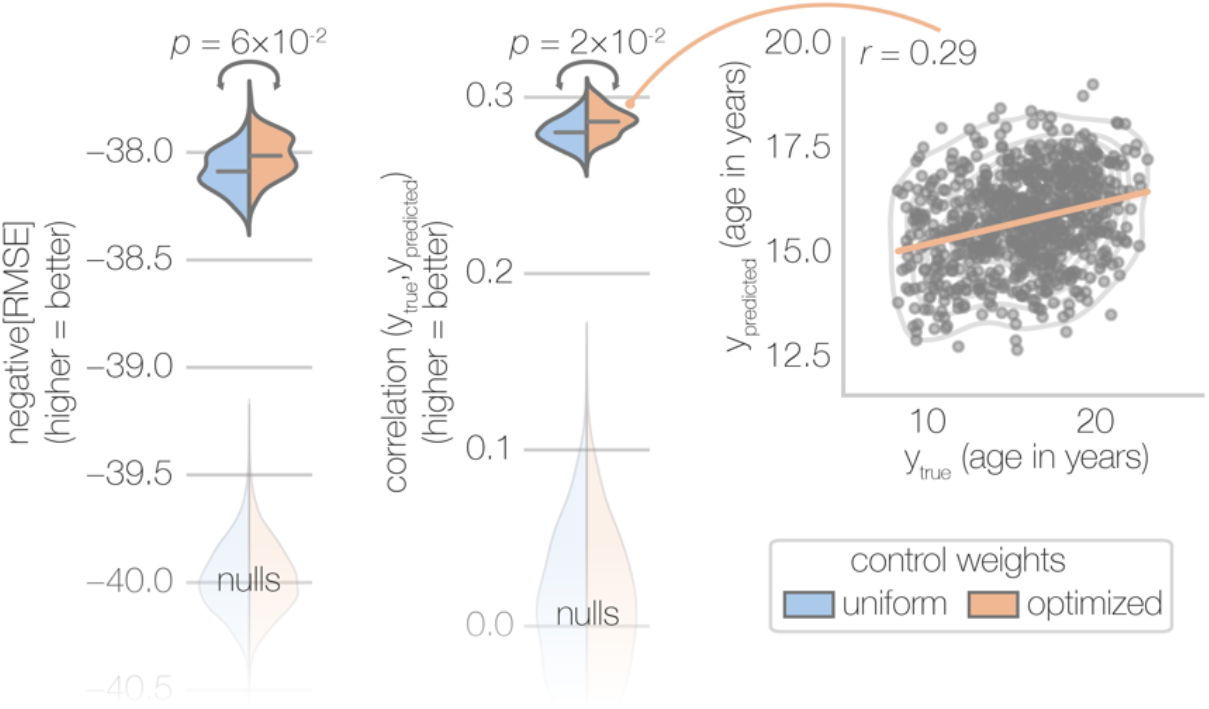
Optimized energy asymmetries improve the out-of-sample prediction of participants’ age. A cross-validated out-of-sample regression model revealed that optimized energy asymmetries (peach) better predicted participants’ age compared to uniform control weights (blue). Out-of-sample prediction was scored using both negative root mean square error (left) and the correlation between true and predicted *y* (right). For the latter, mean prediction performance (gray horizontal lines) was significantly higher for optimized compared to uniformly weighted energy asymmetries (*p*=2×10^−2^). For the former, mean prediction performance was higher for optimized compared to uniformly weighted energy asymmetries but this effect was only marginally significant (*p*=6×10^−2^). Additionally, all mean prediction performance estimates were significantly higher than expected under their respective empirical nulls.

**Figure S5.**
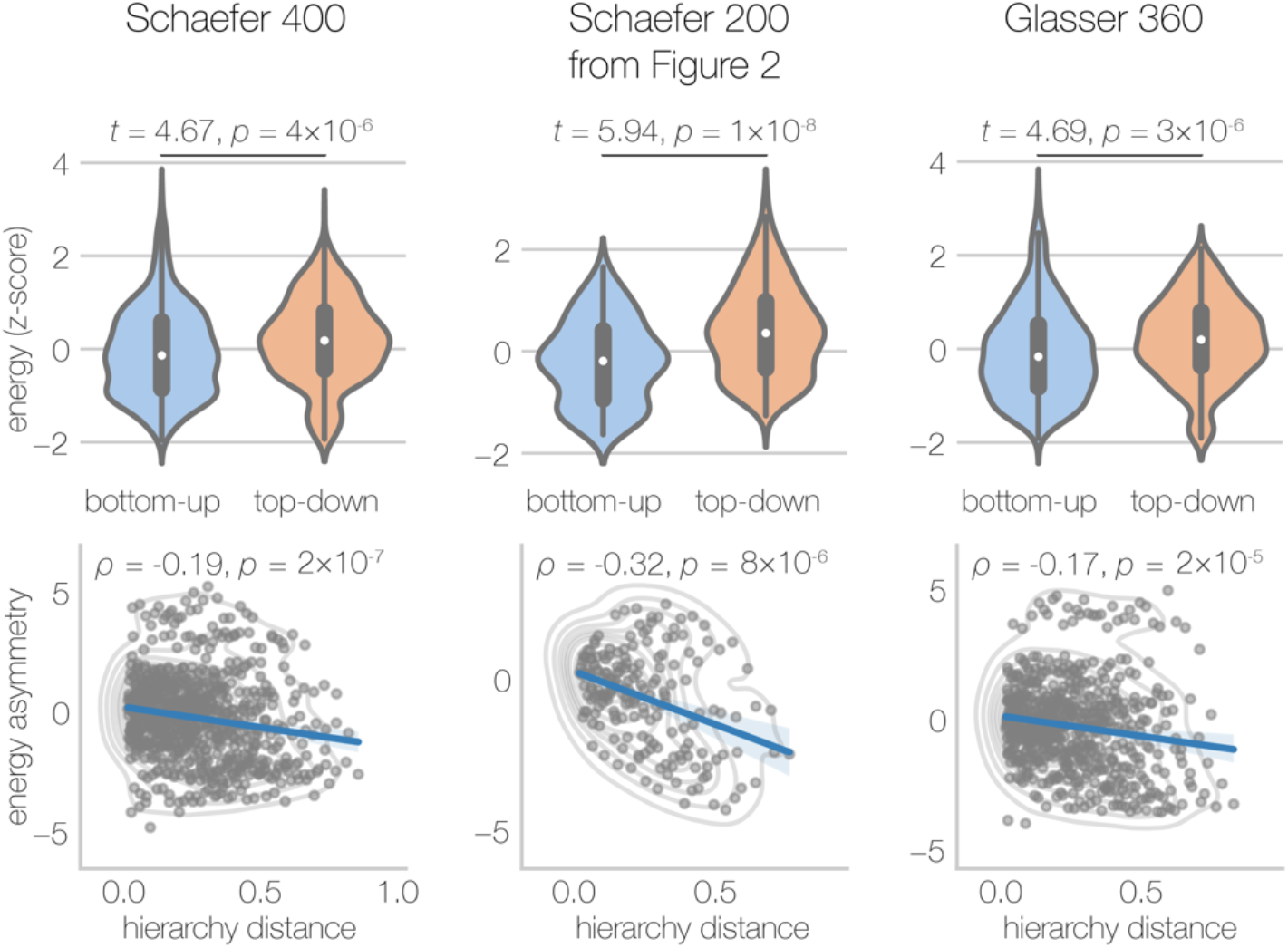
Sensitivity analysis, connectome parcellation. In the main text, we reported network control theory results that were modeled on a connectome built from a parcellation with 200 regions (Schaefer et al., 2018). Here, we examined whether our primary findings were robust to our choice of parcellation by reproducing results from Figure 2 twice, once using a higher resolution version of the same parcellation (Schaefer 400, left) and once using a parcellation with 360 regions defined according to different criteria (Glasser 360, right) (Glasser et al., 2016). For both Schaefer 400 and Glasser 360, we observed that bottom-up energy was significantly lower than top-down energy and that hierarchical distance correlated negatively with energy asymmetry. However, we found that the distance correlation was weaker for both Schaefer 400 and Glasser 360 (compared to the original parcellation, Schaefer 200), suggesting that this effect may be somewhat scale/parcellation dependent.

**Figure S6.**
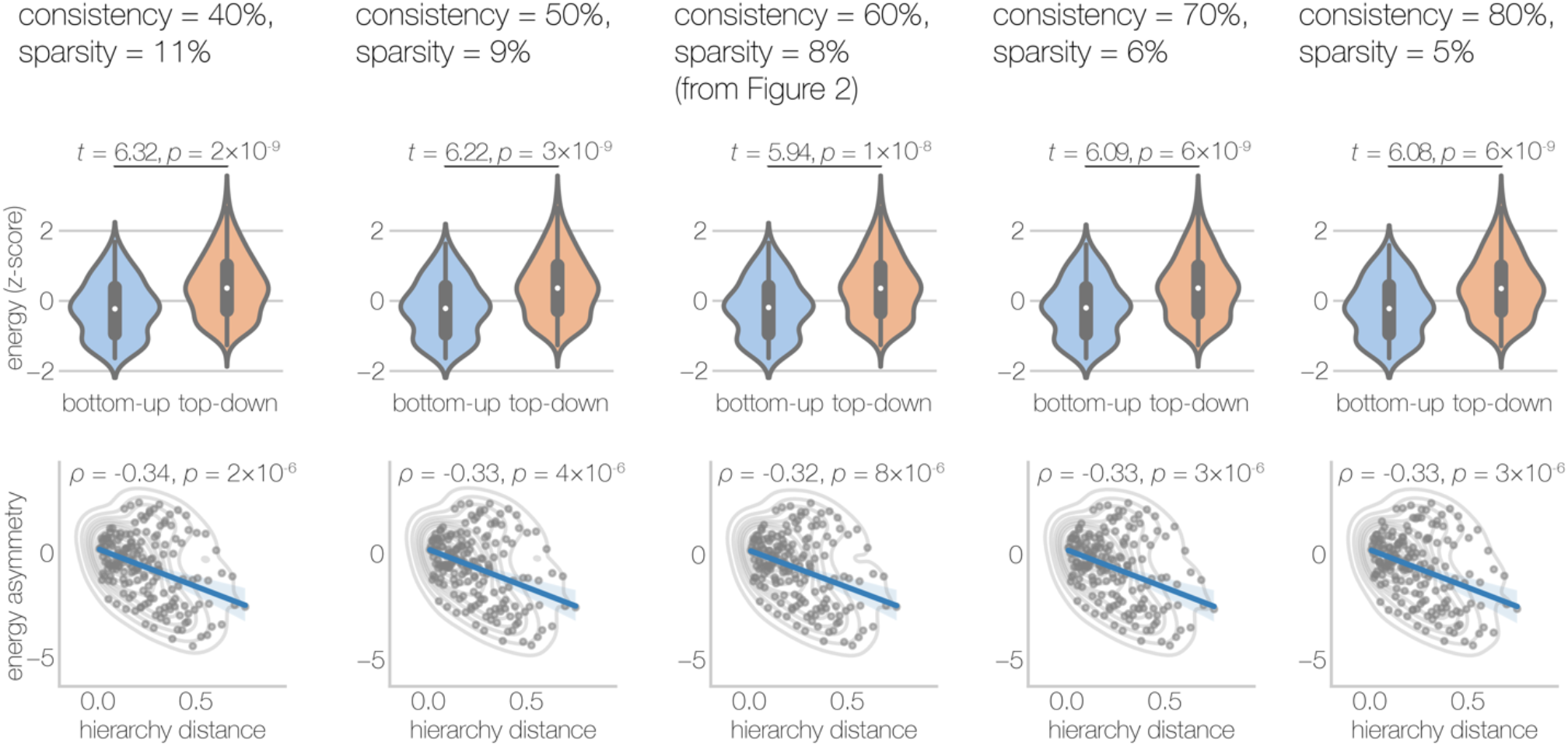
Sensitivity analysis, connectome sparsity. In the main text, we reported network control theory results derived from a group-averaged connectome that was thresholded to retain edges that were present in at least 60% of participants. This thresholding yielded a connectome with 8% sparsity. Here, we examined whether our primary findings were robust to that choice by reproducing results from Figure 2 at a range of consistency thresholds (40%, 50%, 70%, and 80%). We observed that our results were highly consistent across this range of thresholds.

**Figure S7.**
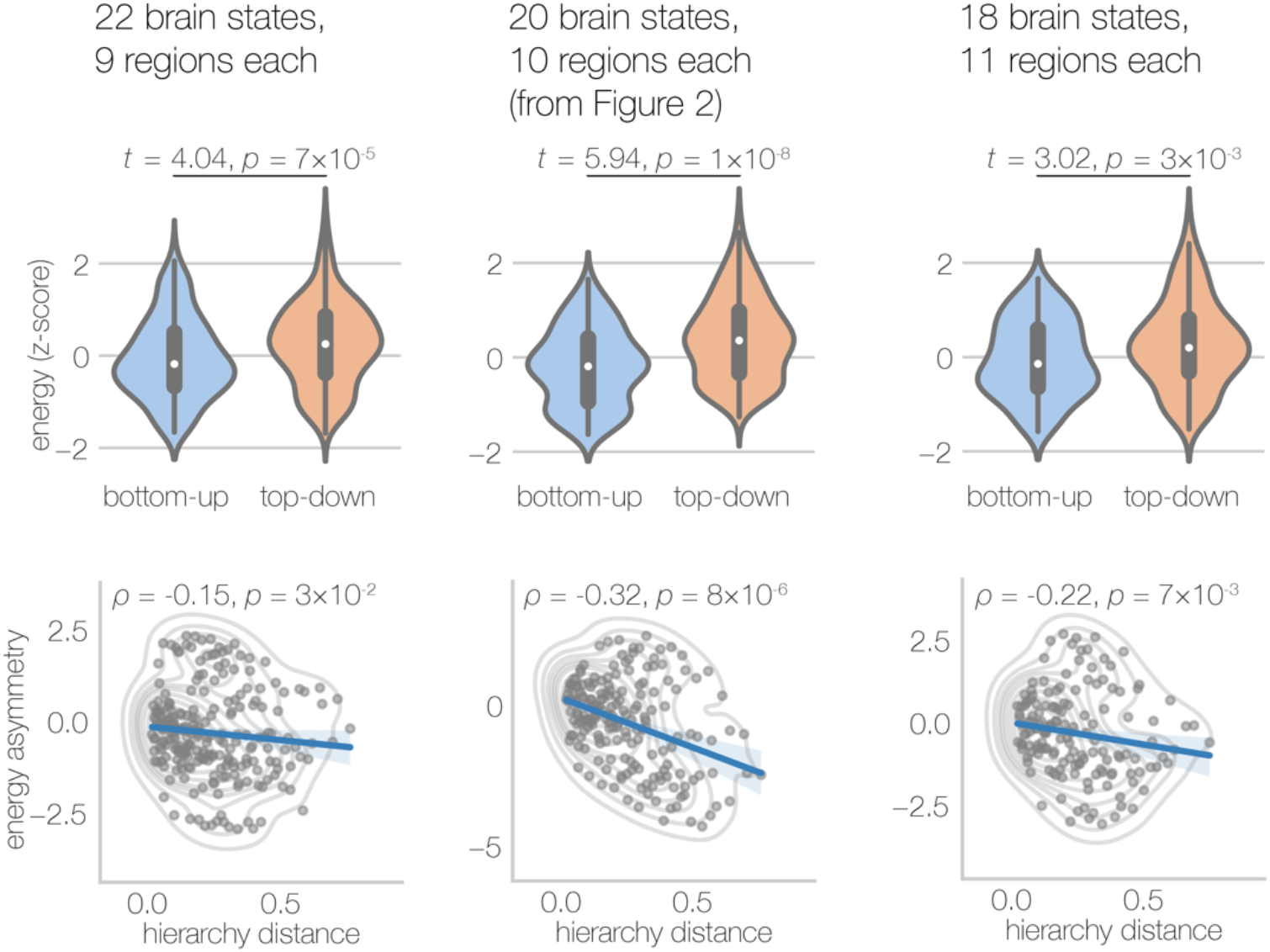
Sensitivity analysis, number of regions per brain state. In the main text, we reported network control theory results for transitions between 20 cytoarchitectonic brain states, each comprising 10 regions. Here, we examined whether our primary findings were robust to that choice by reproducing results from Figure 2 twice, once incrementing and once decrementing the size of brain states by one region. For both brain states of sizes 9 and 11, we observed that bottom-up energy was significantly lower than top-down energy and that hierarchical distance correlated negatively with energy asymmetry. However, we found that the distance correlation was weaker for state sizes 9 and 11 (compared to the original size of 10), suggesting that this effect may be somewhat scale dependent.

**Figure S8.**
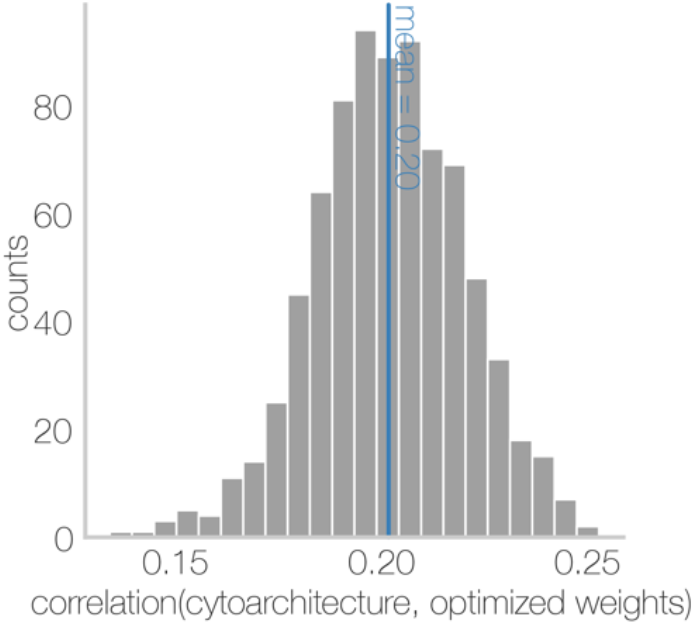
Distributions of correlations between subject-specific optimized weights and the sensory-fugal axis of cytoarchitecture. For each subject, optimized control weights were generated for each state transition (see main text) and correlated with the sensory-fugal axis of cytoarchitecture. Then, for each subject, correlations were averaged over all state transitions yielding a single correlation per subject; these summary correlations are plotted here.

